# SEC16A/MAG5-mediated ER export shapes ABA receptor homeostasis

**DOI:** 10.64898/2026.11.06.761007

**Authors:** José Antonio Berlanga-Torres, Adela Villasante-Fernández, Cecilia Costigliolo-Rojas, Jeanette Moulinier-Anzola, Mar Bono, Pedro L. Rodriguez, Barbara Korbei, María Jesús Marcote, Fernando Aniento, Alejandro Ferrando, Borja Belda-Palazón

**Affiliations:** Instituto de Biología Molecular y Celular de Plantas (IBMCP), CSIC, Universitat Politècnica de València, 46022 Valencia, Spain; Department of Biotechnology and Food Sciences, Institute of Molecular Plant Biology, BOKU University, Muthgasse 18, 1190, Vienna, Austria; Departamento de Bioquímica y Biología Molecular, Facultad de Farmacia, Instituto Universitario de Biotecnología y Biomedicina (BIOTECMED), Universitat de València, 46100, Burjassot, Spain

**Keywords:** ABA receptors, ABA signaling pathway, ER, Golgi, COPII, SEC16A/MAG5, secretory pathway

## Abstract

- The phytohormone abscisic acid (ABA) controls plant adaptation to environmental stress through PYR/PYL/RCAR receptors. Regulated receptor turnover through ubiquitination and endosomal trafficking, including ESCRT-dependent vacuolar degradation, is a major mechanism controlling ABA signalling. However, whether ABA receptor homeostasis is established during the early secretory pathway remains unknown.
- Using complementary cell biological and biochemical approaches, we define a constitutive early secretory pool of the ABA receptors PYR1 and PYL4 associated with the endoplasmic reticulum (ER) and the Golgi apparatus (GA).
- Genetic, cell biological and pharmacological analyses reveal that the COPII scaffold protein SEC16A (also known as MAG5) is required for the proper trafficking and homeostasis of this receptor pool. Loss of MAG5 causes intracellular receptor accumulation, impaired trafficking towards late endosomal compartments and enhanced ABA signalling without affecting endogenous ABA accumulation.
- Together, our findings establish the early secretory pathway as an integral component of the ABA signalling network and demonstrate that MAG5-mediated ER export shapes ABA receptor homeostasis by coordinating receptor trafficking throughout the endomembrane system, revealing the ER as a key regulatory compartment for ABA receptor homeostasis.

## Introduction

The endomembrane system is a highly dynamic network that coordinates intracellular protein trafficking, membrane identity and cellular signalling, thereby enabling plants to continuously adapt their growth and development to changing environmental conditions. Rather than functioning as a constitutive transport route, the secretory pathway is increasingly recognized as a regulatory platform whose organization is remodelled according to developmental stage, secretory demand and environmental stress. Such plasticity not only maintains membrane homeostasis but also determines the abundance, localization and activity of signalling proteins, placing protein trafficking alongside transcriptional and translational regulation as a fundamental mechanism controlling cellular responses (Paez Valencia *et al*., 2016; Wang *et al*., 2020; Aniento *et al*., 2022).

Among the signalling pathways that require particularly tight spatial regulation, abscisic acid (ABA) signalling plays a central role in orchestrating plant adaptation to drought, salinity, osmotic stress and other adverse environmental conditions. Perception of ABA by members of the PYRABACTIN RESISTANCE/PYR1-LIKE/REGULATORY COMPONENT OF ABA RECEPTOR (PYR/PYL/RCAR) family initiates a signalling cascade in which clade A PROTEIN PHOSPHATASE 2Cs (PP2Cs) are inhibited, allowing activation of SNF1-RELATED PROTEIN KINASE 2 (SnRK2) kinases and downstream transcriptional and physiological responses (Cutler *et al*., 2010; Yoshida *et al*., 2019). Because ABA signalling controls essential processes including stomatal closure, seed dormancy, germination and stress acclimation, the abundance, subcellular localization and activity of ABA receptors must be tightly coordinated in space and time to ensure signalling outputs that are both robust and rapidly reversible. Consequently, the compartmentalization of ABA receptors has emerged as a central determinant of signalling intensity, duration and sensitivity (Rodriguez *et al*., 2019; Coego *et al*., 2021; Qin *et al*., 2026).

Consistent with this concept, ABA receptor homeostasis is regulated through multiple sequential layers acting across the endomembrane system. In addition to transcriptional regulation, receptor abundance is extensively controlled by ubiquitination and selective protein degradation. Several E3 ubiquitin ligases operate at distinct cellular compartments, including CRL4 complexes, RBR-type RFA ligases and the plasma membrane-associated RING E3 ligase RSL1, promoting proteasomal or vacuolar degradation depending on receptor localization (Irigoyen *et al*., 2014; Bueso *et al*., 2014; Fernandez *et al*., 2020; Coego *et al*., 2021). However, receptor proteostasis is not solely determined by protein turnover. Members of the C2-domain ABA-related (CAR) protein family recruit specific PYR/PYL receptors to the plasma membrane in a calcium-dependent manner, thereby facilitating efficient ABA perception by controlling receptor positioning (Rodriguez *et al*., 2015; Diaz *et al*., 2016). Importantly, ABA signalling also regulates membrane-delimited processes, including the activity of plasma membrane ion channels that control ion and water fluxes during stomatal closure. The spatial organization of ABA signalling components at or near the plasma membrane may therefore directly influence the efficiency and specificity of physiological ABA responses (Geiger *et al*., 2009; Imes *et al*., 2013; Kollist *et al*., 2014). More recently, the endoplasmic reticulum (ER)-associated E3 ubiquitin ligase DOA10A was shown to mono-ubiquitinate ABA receptors, promoting their trafficking through the endomembrane system towards the plasma membrane without affecting receptor stability (Liu *et al*., 2025). This finding fundamentally expanded the role of ubiquitination in ABA signalling, demonstrating that ubiquitin modifications regulate receptor localization independently of protein degradation. Following receptor activation and internalization, components of the ESCRT machinery, including FYVE1/FREE1, FYVE4, VPS23A, ALIX and TOM1-LIKE (TOL) proteins, mediate endosomal sorting and vacuolar degradation of ABA receptors (Belda-Palazon *et al*., 2016; Yu *et al*., 2016; García-León *et al*., 2019; Moulinier-Anzola *et al*., 2024; Chen *et al*., 2025). Together, these studies reveal that receptor abundance and subcellular distribution depend on the coordinated regulation of intracellular trafficking across the endomembrane system.

Despite these advances, the spatial distribution of ABA receptors within the early secretory pathway remains poorly understood. Export of newly synthesized membrane and secretory proteins from the ER occurs at specialized ER exit sites (ERES) through the assembly of COPII-coated carriers (Brandizzi & Barlowe, 2013). This process is initiated by activation of the small GTPase SAR1 by the ER-localized guanine nucleotide exchange factor SEC12, promoting SAR1 membrane insertion and recruitment of the inner coat complex SEC23/SEC24, which mediates cargo selection. Assembly of the outer coat complex SEC13/SEC31 subsequently drives COPII carrier formation and ER-to-Golgi transport, whereas cycles of SAR1 GTP binding and hydrolysis coordinate coat assembly, carrier scission and coat disassembly. Consequently, modulation of COPII dynamics enables cells to rapidly adjust secretory flux and protein distribution in response to developmental and environmental cues (Brandizzi, 2018; Aniento *et al*., 2022)

Supporting this view, increasing evidence indicates that the architecture and activity of the early secretory pathway is actively remodelled during environmental adaptation (Brandizzi, 2018; Aniento *et al*., 2022). A recent study demonstrated that ABA remodels COPII dynamics and early secretory trafficking, revealing that hormone signalling directly targets the machinery responsible for ER export (Li *et al*., 2021). These observations suggest that modulation of ER export constitutes an additional regulatory layer controlling signalling specificity by regulating the availability of selected signalling proteins before they enter downstream trafficking pathways.

A central organizer of ER export is SEC16, a large peripheral membrane protein that plays a unique role in COPII assembly because it is not a structural component of the coat itself but instead acts as a master scaffold that spatially organizes ER exit sites. SEC16 orchestrates the spatial organization and stability of ER exit sites by coordinating COPII assembly and turnover, and this role is broadly conserved across eukaryotes (Connerly *et al*., 2005; Watson *et al*., 2006; Brandizzi & Barlowe, 2013). In *Arabidopsis thaliana*, two closely related SEC16 homologues, SEC16A (also known as MAG5) and SEC16B, have been identified. Although both proteins share high sequence similarity, MAG5 is the predominant isoform required for ER exit site organization and efficient ER-to-Golgi trafficking, whereas SEC16B appears to play a more limited or partially redundant role in early secretory trafficking (Takagi *et al*., 2013). Given that ABA signalling remodels the early secretory pathway through the regulation of COPII-dependent trafficking (Li *et al*., 2021), MAG5 represents an attractive candidate for coupling hormone signalling with early secretory trafficking.

Here, we demonstrate that the early secretory pathway constitutes a previously unrecognized regulatory layer controlling ABA receptor homeostasis. By combining cell biology, genetic and biochemical approaches, we show that MAG5 regulates the homeostasis of a distinct ER-Golgi-associated receptor pool, whose abundance and subcellular organization depend on functional ER export. Disruption of MAG5 alters receptor distribution and abundance and enhances ABA sensitivity, revealing that early secretory trafficking actively shapes hormone perception rather than merely supporting constitutive protein secretion. Together, our findings establish the ER–Golgi interface as an active signalling hub that integrates membrane trafficking with hormone perception, expanding current models of ABA signalling during plant stress adaptation.

## Materials and Methods

### Plant materials and growth conditions

*Arabidopsis thaliana* ecotype Columbia-0 (Col-0) was used as the wild-type control. All mutant and transgenic lines used in this study are listed in Table **S1**. Unless otherwise indicated, plants were grown under long-day conditions (16 h light/8 h dark; 22°C/18°C) with a light intensity of 100-120 μmol m⁻² s⁻¹. Surface-sterilized seeds were sown on half-strength Murashige and Skoog (½ MS) medium with vitamins and MES (Duchefa M0255) supplemented with 1.03 g/L of MES and 0.8% (w/v) phytoagar (pH 5.7). Plates were sealed with Micropore™ tape, stratified in darkness at 4°C for 2 d, and then transferred to the growth chamber.

### Molecular cloning and plasmid construction

Gateway® cloning technology (Invitrogen) was used for the generation of all constructs unless otherwise indicated. PCR-amplified coding sequences were first cloned into the Gateway entry vector pDONR™/Zeo by BP recombination and subsequently transferred into the appropriate Gateway destination vectors by LR recombination according to the manufacturer’s instructions. All constructs were verified by sequencing. A complete list of primers, plasmids and constructs used in this study is provided in Tables **S2** and **S3**.

### *Arabidopsis* transformation

Stable transgenic *A. thaliana* lines were generated by *Agrobacterium tumefaciens*-mediated transformation of flowering Col-0 plants using the floral dip method (Clough & Bent, 1998). T2 progeny were screened to identify lines carrying a single transgene insertion, and homozygous T3 lines were used for subsequent analyses. All transgenic lines used in this study are listed in Table **S1**.

### ABA and salt sensitivity assays during germination and early seedling development

Surface-sterilized seeds were sown on ½ MS medium supplemented with 0.3 or 0.5 µM ABA, and 100 or 150 mM NaCl. Following stratification, plates were transferred to long-day growth conditions and seed germination, and early seedling development were monitored over time. Germination was scored as radicle emergence 48-72 h after transfer to the growth chamber, whereas early seedling establishment was quantified 72-96 h after transfer as the percentage of seedlings developing fully expanded green cotyledons. For each genotype and treatment, at least 50 seeds were analysed per biological replicate.

### Primary root growth assays

For primary root growth assays, surface-sterilized seeds were germinated on ½ MS medium and grown for 5 d under long-day conditions. Seedlings were then transferred to fresh ½ MS plates supplemented with 5 µM of ABA, and allowed to grow for an additional 8 d. At the end of the treatment, seedlings were subsequently transferred to fresh agar plates, carefully straightened and photographed. Primary root growth was quantified from the scanned images using Fiji/ImageJ (Schindelin *et al*., 2012).

### ABA quantification

Endogenous ABA was quantified in 6-day-old seedlings. Approximately 100 mg of fresh tissue was extracted with methanol/water/acetic acid (80:19:1, v/v/v) containing deuterium-labelled ABA as an internal standard. Following centrifugation, the supernatant was dried under vacuum, resuspended in 1% (v/v) acetic acid and purified by reverse-phase solid-phase extraction. The eluate was dried, resuspended in 5% (v/v) acetonitrile containing 1% (v/v) acetic acid, and analysed by ultra-high-performance liquid chromatography coupled to high-resolution mass spectrometry (UHPLC-HRMS). ABA was quantified using a Q Exactive Orbitrap mass spectrometer (Thermo Fisher Scientific, Waltham, MA, USA) operated in selected ion monitoring (SIM) mode at the Hormone Quantification Service of the IBMCP (Valencia, Spain).

### Infrared thermography and stomatal aperture measurements

Infrared thermography was performed on 6-week-old *A. thaliana* plants grown under short-day conditions (8 h light/16 h dark, 21°C/19°C, 100 μmol m⁻² s⁻¹). Plants were sprayed with 20 mM MES (pH 5.7) containing 0.02% (v/v) Silwet L-77 supplemented with the indicated concentration of ABA or an equivalent volume of DMSO as a mock control. Thermal images were acquired 3 h after treatment using an FLIR E95 infrared camera (FLIR Systems) equipped with a 42° lens. Images were analysed using FLIR Tools software. Mean leaf temperature was determined from 15 independent regions of interest distributed across 4-6 rosette leaves per plant. At least six plants were analysed per genotype in each experiment, and experiments were performed independently at least twice.

Stomatal aperture was measured in cotyledons of 8-day-old seedlings. Seedlings were harvested directly into 95% ethanol after 2–3 h of light exposure and fixed for 3–4 h. Samples were subsequently incubated for 10 min each in 70%, 50% and 20% ethanol and cleared overnight in a 1:3:8 glycerol:water:chloral hydrate solution. Cotyledons were mounted with the abaxial surface facing upwards and imaged using a 20× objective. At least one image from a comparable region of the cotyledon was acquired for each of 9 individual seedlings, with at least 20 stomata quantified per seedling. Stomatal parameters were quantified using ImageJ. Stomatal aperture was expressed as the ratio of stomatal pore width to length.

### Transient expression in *Nicotiana benthamiana*

Transient expression assays were performed in fully expanded leaves of 4-5-week-old *Nicotiana benthamiana* plants by *A. tumefaciens*-mediated infiltration as described previously (Belda-Palazón & Rodriguez, 2020, 2022). Briefly, *A. tumefaciens* strains carrying the indicated binary constructs were grown overnight, collected by centrifugation and resuspended in infiltration buffer (10 mM MgCl₂, 10 mM MES, pH 5.6, and 150 μM acetosyringone) to an OD_600_ of 2.0. Bacterial suspensions were incubated at room temperature for 3-5 h before infiltration. Immediately before infiltration, cultures carrying the desired constructs were mixed and diluted in infiltration buffer to the final OD_600_ required for each experiment, resulting in a final OD_600_ of 0.25-0.5 for each individual construct.

For co-expression and colocalization experiments, the indicated fluorescent protein fusions were co-expressed with the corresponding organelle markers. For bimolecular fluorescence complementation (BiFC) assays, *A. tumefaciens* cultures carrying the indicated YFP^N^ and YFP^C^ fusion constructs were co-infiltrated together with fluorescent subcellular markers (Belda-Palazón *et al*., 2012). Leaves were analysed by confocal laser scanning microscopy 72-96 h after infiltration. A complete list of constructs used is provided in Table **S2**.

### Pharmacological treatments and fluorescent probes

For pharmacological treatments, 5-day-old *A. thaliana* seedlings expressing the indicated fluorescent reporters in the corresponding genetic backgrounds were incubated in liquid ½ MS medium supplemented with the indicated compounds or an equivalent volume of DMSO (mock) for 1 h. Brefeldin A (BFA; TargetMol) was used at 50 μM and wortmannin (WM; TargetMol) at 33 μM.

For endocytic labelling, seedlings were incubated with FM4-64 (2 μM; Sigma-Aldrich) for 10 min following BFA treatment. To visualize late acidic endosomal/vacuolar compartments, following WM treatment seedlings were incubated with LysoTracker™ Deep Red (1 μM; Thermo Fisher Scientific) for an additional 10 min before confocal imaging. All treatments were performed immediately prior to confocal microscopy.

### Confocal laser scanning microscopy

Confocal imaging was performed using a Stellaris 8 laser scanning confocal microscope (Leica Microsystems) equipped with a HC PL APO 40×/1.30 OIL objective and HyD detectors. GFP/YFP fluorescence was excited at 488 nm and detected between 500 and 530 nm, whereas RFP/mCherry fluorescence was excited at 561 nm and detected between 600 and 630 nm. FM4-64 was excited at 488 nm and detected between 610 and 630 nm, whereas LysoTracker™ Deep Red was excited at 647 nm and detected between 660 and 680 nm. For multicolour imaging, fluorophores were acquired sequentially to avoid spectral overlap. Pinhole size was adjusted to 1 Airy unit for each channel. Images were acquired using Leica LAS X software and processed using Fiji/ImageJ.

### Subcellular fractionation and sucrose density gradient centrifugation

Microsomal membrane fractions were prepared from two-week-old wild-type (Col-0) *A. thaliana* seedlings. Approximately 1 g of frozen seedling tissue was ground to a fine powder in liquid nitrogen and homogenized in three volumes of ice-cold homogenization buffer (20 mM Tris-HCl, pH 8.0, 250 mM sucrose, 5% glycerol, 5 mM MgCl₂, 5 mM KCl, 5 mM EDTA and 14 mM β-mercaptoethanol) supplemented with cOmplete™, Mini, EDTA-free Protease Inhibitor Cocktail (Roche). Homogenates were incubated on ice for 30 min with gentle inversion every 5 min, filtered through one layer of Miracloth and centrifuged at 8,000 × g for 10 min at 4°C to remove cell debris and large organelles. The resulting supernatant (S8) was ultracentrifuged at 100,000 × g for 30 min at 4°C to obtain the soluble (S100) and microsomal membrane (P100) fractions. The P8 and P100 pellets were resuspended in homogenization buffer. An aliquot of each fraction was subsequently mixed with 4× Laemmli sample buffer, heated at 70°C for 10 min, cooled on ice, and analysed by SDS–PAGE and immunoblotting.

For density gradient fractionation, the P100 microsomal fraction was layered onto continuous 10-50% (w/v) sucrose gradients prepared in gradient buffer containing 40 mM Tris-HCl (pH 8.5), 20 mM KCl and 10 mM MgCl₂. Gradients were centrifuged overnight at 35,000 rpm in a SW40 Ti rotor (Beckman Coulter) at 4°C without brake. Following centrifugation, gradients were collected manually from the top into sixteen fractions of equal volume (F1-F16). Proteins were precipitated overnight at -20°C by addition of one volume of 20% trichloroacetic acid (TCA), centrifuged at 12,000 × g for 15 min at 4°C, washed twice with cold acetone, air-dried, and resuspended in 2× Laemmli sample buffer. Samples were subsequently heated at 70°C for 10 min, cooled on ice and equal volumes from each fraction were analysed by SDS-PAGE and immunoblotting using antibodies against PYR1 and organelle-specific marker proteins (Table **S4**).

### Protein extraction and immunoblot analysis

For immunoblot analyses, approximately 100 mg of frozen *A. thaliana* tissue was ground to a fine powder in liquid nitrogen and homogenized in two volumes of 2× Laemmli extraction buffer (126 mM Tris-HCl, pH 6.8, 4% SDS, 20% glycerol, 0.05% bromophenol blue and 4% β-mercaptoethanol). Homogenates were incubated on ice for 20-30 min with intermittent vortexing, boiled at 95°C for 10-15 min, cooled on ice and centrifuged at 12,000 × g for 15 min. The resulting supernatants were used as total protein extracts. Equal volumes of protein extracts were separated by SDS-PAGE and transferred onto PVDF membranes.

Membranes were blocked in 5% (w/v) non-fat dry milk in TBS-T and incubated overnight at 4°C with the indicated primary antibodies. Following incubation with horseradish peroxidase-conjugated secondary antibodies, immunoreactive bands were detected using enhanced chemiluminescence (ECL). Band intensities were quantified using Fiji/ImageJ and normalized to the corresponding loading controls. Antibodies used in this study are listed in Table **S4**.

### RNA extraction and RT-qPCR analysis

For gene expression analyses, total RNA was extracted from whole *A. thaliana* seedlings using NZY Total RNA Isolation kit (NZYtech) according to the manufacturer’s instructions. RNA samples were treated with the TURBO DNA-free™ Kit (Invitrogen, Thermo Fisher Scientific). RNA concentration and purity were determined using a NanoDrop™ One spectrophotometer (Thermo Fisher Scientific). First-strand cDNA was synthesized from 1 μg of total RNA using the PrimeScript™ RT reagent Kit (Takara Bio Inc.) following the manufacturer’s instructions.

Quantitative real-time PCR (RT-qPCR) was performed using HOT FIREPol® EvaGreen® qPCR Mix Plus (Solis Biodyne) in a final reaction volume of 10 μL containing 0.4 μL of cDNA and 500 nM of each primer. Reactions were run on a QuantStudio™ 3 Real-Time PCR System (Applied Biosystems). Relative transcript abundance was calculated using the 2^−ΔCt^, with *PDF2* used as the internal reference gene. Each biological replicate was analysed in three technical replicates. Primer sequences are listed in Table **S3**.

### Statistical analyses

Statistical analyses were performed using GraphPad Prism 8 (GraphPad Software). Data are presented as mean ± SEM. Statistical significance was assessed by one-way or two-way ANOVA followed by Tukey’s or Dunnett’s test multiple comparison test, or by Student’s t-test when appropriate. Details of the statistical tests, sample sizes and significance levels are provided in the corresponding figure legends.

## Results

### PYR1 and PYL4 constitutively occupy a previously unrecognized ER-Golgi-associated subpopulation

A recent study demonstrated that mono-ubiquitination promotes ABA receptor trafficking towards the plasma membrane, suggesting that receptor localization may already be regulated during early secretory trafficking (Liu *et al*., 2025). Whether ABA receptors constitutively occupy a distinct ER-Golgi-associated subpopulation before undergoing this regulated trafficking, however, has remained unknown. To address this question, we examined the subcellular distribution of functional PYR1 and PYL4 reporters using complementary imaging and biochemical approaches.

To investigate whether ABA receptors are associated with the early secretory pathway, we first analysed the subcellular distribution of GFP-PYR1 and GFP-PYL4 translational fusions by confocal microscopy in *N. benthamiana* using markers of distinct endomembrane compartments (Nelson *et al*., 2007; Serra-Soriano *et al*., 2014; Bernat-Silvestre *et al*., 2022). In agreement with previous reports, both receptors displayed predominant nucleocytoplasmic localization together with fluorescence at the cell periphery (Belda-Palazon *et al*., 2016). In addition, both receptors consistently accumulated in a network-like reticular pattern and in discrete intracellular puncta (Figs **1a,b**). The reticular signal overlapped with the ER luminal marker ChFP-KDEL, whereas a subset of the intracellular puncta colocalized with the cis-Golgi marker ManI-RFP, indicating that both PYR1 and PYL4 are present within the early secretory pathway. Time-lapse confocal imaging further showed that the ER-associated PYR1 signal remained dynamically associated with the ER network during its continuous remodelling (Video/Movie **S1**).

**Figure 1.**
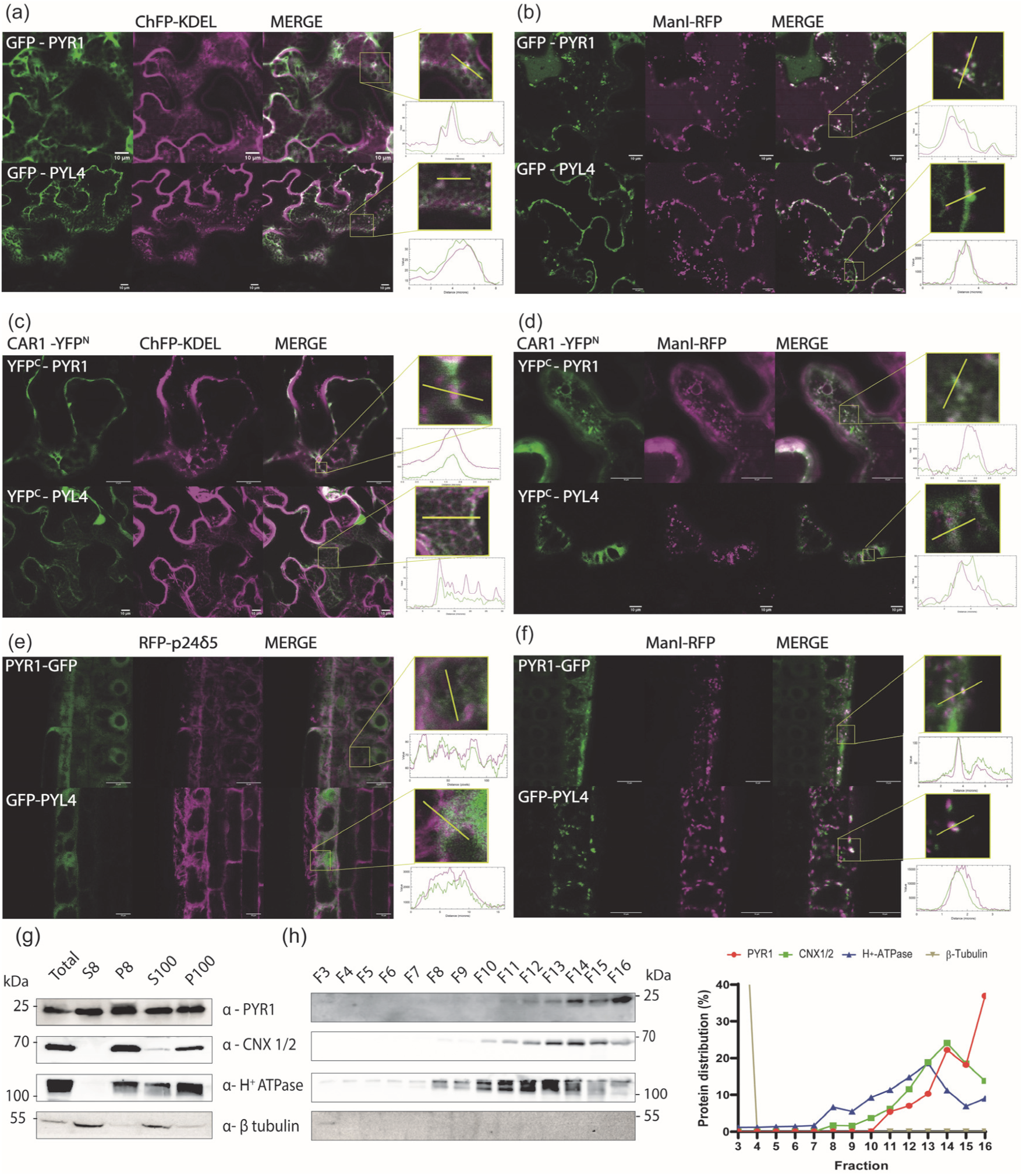
PYR1 and PYL4 define a constitutive receptor pool associated with the early secretory pathway. (a, b) Representative confocal images of *N. benthamiana* epidermal cells transiently expressing GFP-PYR1 or GFP-PYL4 together with the ER marker ChFP-KDEL (a) or the cis-Golgi marker ManI-RFP (b). (c, d) Representative BiFC confocal images following transient co-expression in *N. benthamiana* epidermal cells of CAR1-YFP^N^ and YFP^C^-PYR1 or YFP^C^-PYL4 together with ChFP-KDEL(c) and ManI-RFP (d). (e, f) Confocal images of root epidermal cells from transgenic *A. thaliana* seedlings expressing *pPYR1::PYR1-GFP* or *35S::GFP-PYL4* together with the ER marker RFP-p24δ5 (e), or the cis-Golgi marker ManI-RFP (f). Insets show magnified regions indicated by the yellow boxes. Fluorescence intensity profiles along the yellow lines are also shown on the right. Scale bars: 10 μm. (g) Immunoblot analyses of protein extracts from wild-type (Col-0) seedlings after subcellar fractionation, using the indicated antibodies. Total protein extract (Total), soluble fraction after centrifugation at 8,000 × g (S8), pellet after centrifugation at 8,000 × g (P8), soluble supernatant after ultracentrifugation at 100,000 × g (S100) and microsomal membrane pellet after ultracentrifugation at 100,000 × g (P100). CNX1/2, H^+^-ATPase and β-tubulin were used as markers for the ER, plasma membrane and cytosolic fractions, respectively. (h) Immunoblot analyses of protein extracts after continuous sucrose density gradient fractionation of the P100 microsomal fraction, using the indicated antibodies. The corresponding densitometric analysis is shown on the right.

Since PYR1 and PYL4 are soluble proteins lacking transmembrane domains, their association with the ER-Golgi endomembrane system prompted us to investigate whether receptor interactions with membrane-associated proteins occur at these compartments. CAR proteins are peripheral membrane proteins and well-established interactors of ABA receptors (Rodriguez *et al*., 2015; Diaz *et al*., 2016; Cui *et al*., 2023). We therefore analysed the subcellular localization of PYR1-CAR1 and PYL4-CAR1 complexes by BiFC. BiFC fluorescence was detected at the cell periphery, as previously reported (Rodriguez *et al*., 2015; Diaz *et al*., 2016), but also in a reticular network and in discrete intracellular puncta (Fig. **1c,d**). Co-expression with the ER marker KDEL revealed extensive overlap of the reticular BiFC signal with the ER network, whereas co-expression with the cis-Golgi marker ManI showed that the punctate BiFC structures corresponded to Golgi-associated compartments. Line-scan intensity profiles further supported the spatial coincidence of the BiFC signal with both ER and cis-Golgi markers (Fig. **1c,d**). Moreover, disruption of the membrane-binding capacity of CAR1 by the DADA mutations largely abolished the ER- and Golgi-associated BiFC pattern (Fig. **S1**) (Diaz *et al*., 2016), indicating that membrane association of CAR1 is required for localization of the receptor-CAR complexes within the early secretory pathway.

To validate these observations under native conditions, we next analysed the subcellular localization of PYR1 and PYL4 in transgenic *A. thaliana* lines. Importantly, PYR1 localization was examined using a functional genomic *pPYR1::PYR1-GFP* reporter expressed under the control of its native promoter and endogenous regulatory sequences (Moulinier-Anzola *et al*., 2024), thereby minimizing potential artefacts associated with transient heterologous expression or constitutive overexpression, and providing an independent validation of the observations obtained by transient expression in *N. benthamiana*. Consistent with the observations in *N. benthamiana*, PYR1-GFP displayed nucleocytoplasmic fluorescence together with a reticular network and discrete intracellular puncta. Co-localization with the ER membrane marker RFP-p24δ5 (Bernat-Silvestre et al., 2020) revealed that the reticular PYR1 signal was associated with the ER network, whereas co-expression with the cis-Golgi marker ManI-RFP showed that a subset of PYR1-positive puncta corresponded to Golgi-associated compartments. GFP-PYL4 exhibited a similar subcellular distribution, indicating that association with the early secretory pathway is a common feature of distinct ABA receptors (Fig. **1e,f**, **S2**).

To independently validate these observations, total protein extracts from wild-type (Col-0) seedlings were subjected to differential centrifugation to isolate a microsomal membrane fraction (P100), which was subsequently resolved by continuous sucrose density gradient centrifugation (Fig. **1g,h**). Differential centrifugation efficiently separated soluble and membrane fractions, as indicated by the enrichment of the ER marker CNX1/2 and the plasma membrane marker H+-ATPase in the P100 fraction, whereas β-tubulin was predominantly recovered in the soluble S100 fraction. Consistent with its partial membrane association, PYR1 was detected in both soluble and microsomal fractions, indicating that a substantial fraction of the protein is membrane-associated (Fig. **1g**). Within the sucrose gradient, PYR1 was broadly distributed across membrane fractions, with its abundance progressively increasing from fraction F11 onwards. Notably, PYR1 showed substantial co-fractionation with the ER marker CNX1/2 across the ER-enriched region (F12-F16), exhibiting a first accumulation peak in fraction F14 that closely overlapped with the maximum abundance of CNX1/2, followed by a second and more pronounced peak in the densest fraction (F16) (Fig. **1h**). In contrast, the plasma membrane marker H+-ATPase displayed a distinct distribution, reaching maximal abundance in lighter fractions, whereas residual β-tubulin was detected only in fraction F3, confirming the efficient enrichment of membrane compartments. Together, these biochemical analyses corroborate microscopy observations and provide independent evidence for the existence of a constitutive ER-associated membrane pool of PYR1 consistent with its localization within the early secretory pathway.

Collectively, these results identify a previously unrecognized constitutive ER–Golgi-associated subpopulation of PYR1 and PYL4, establishing the early secretory pathway as an active regulatory compartment controlling ABA receptor homeostasis.

### MAG5 is required for the normal endomembrane distribution and trafficking of ABA receptors

The identification of a constitutive ER-Golgi-associated subpopulation of ABA receptors prompted us to investigate its relationship with the early secretory trafficking machinery. MAG5 is a core organizer of ER exit sites that defines specialized ER domains where COPII-mediated transport is initiated. To examine their spatial relationship, we analysed *A. thaliana* plants co-expressing the functional ER exit site marker *pMAG5::MAG5-GFP* together with RFP-PYR1 (Takagi *et al*., 2013; Moulinier-Anzola *et al*., 2024).

Confocal microscopy revealed that MAG5 accumulated in discrete punctate structures corresponding to ER exit sites, whereas PYR1 exhibited a broader intracellular distribution. Although a substantial proportion was detected outside MAG5-positive puncta, a subset of receptor-containing puncta closely associated with MAG5 (Fig. **2a**), consistent with the transient recruitment of a fraction of the receptor population to ER exit sites during secretory trafficking. A similar spatial association between MAG5 and PYR1 was also observed following transient co-expression in *N. benthamiana* leaf epidermal cells (Fig. **S3a**), indicating that this spatial relationship is reproducible in an independent heterologous expression system. Moreover, BiFC complexes formed by CAR1 and PYR1 remained closely associated with RFP–MAG5-positive compartments (Fig. **S3b**), and time-lapse confocal imaging showed that this spatial association was maintained during the dynamic movement of these structures (Video/Movie **S2**). Together, these observations further support the close association of ABA receptors with MAG5-positive early secretory compartments.

**Figure 2.**
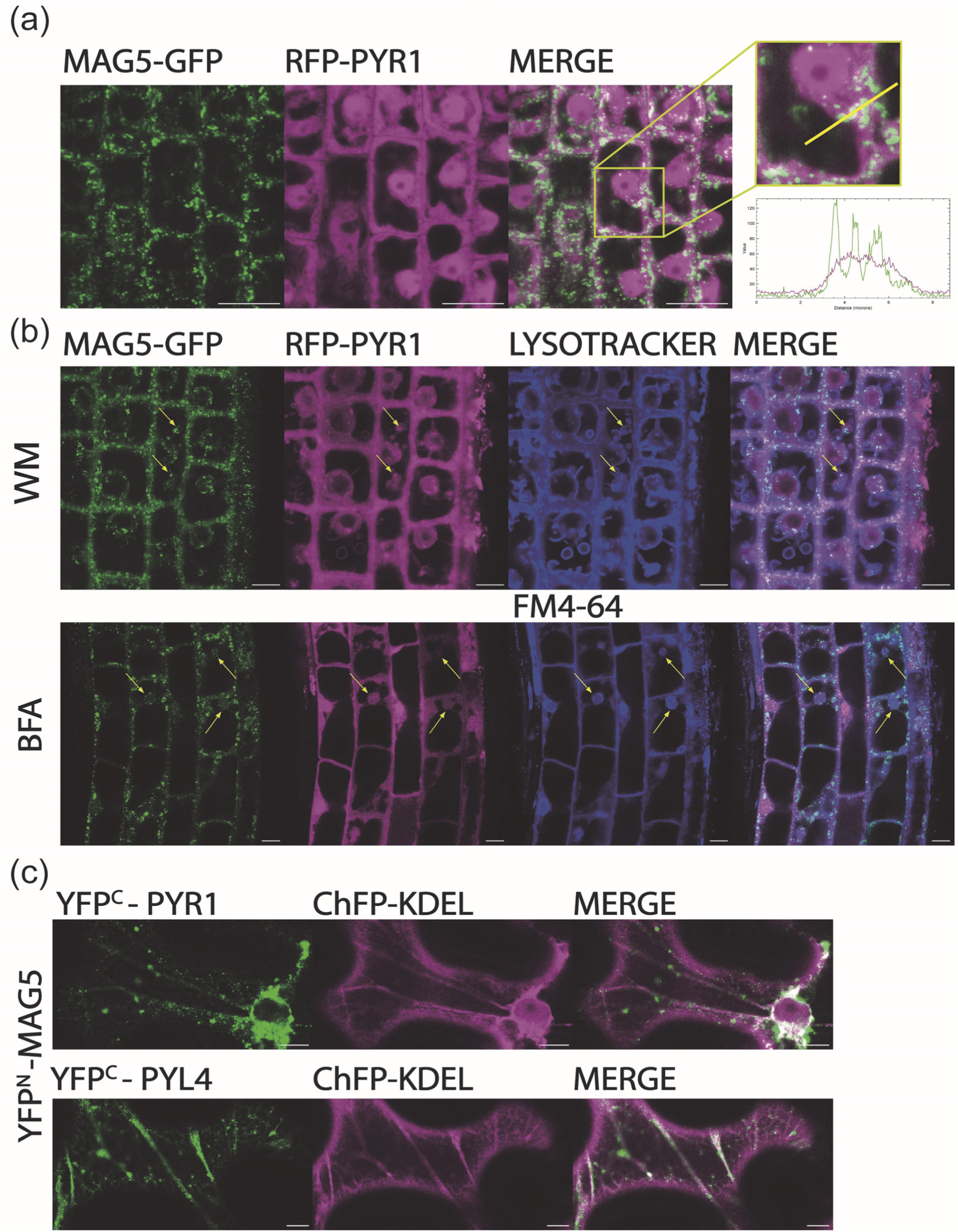
MAG5 associates with ABA receptors within the early secretory pathway. (a) Confocal images of root epidermal cells from transgenic *A. thaliana* seedlings co-expressing *pMAG5::MAG5-GFP* and *35S::RFP-PYR1*. Insets show magnified regions indicated by the yellow boxes. Fluorescence intensity profiles along the yellow line are shown on the right. (b) Subcellular distribution of MAG5-GFP and RFP-PYR1 in root epidermal cells following pharmacological treatments. Five-day-old seedlings were treated with 33 μM WM for 1 h and stained with LysoTracker™ Deep Red, or with 50 μM BFA for 1 h followed by FM4-64 staining. Arrows indicate representative compartments containing both MAG5-GFP and RFP-PYR1 within WM-induced late endosomal structures or BFA bodies. (c) Representative BiFC images following transient co-expression of YFP^N^-MAG5 and YFP^C^-PYR1 or YFP^C^-PYL4 together with ChFP-KDEL in *N. benthamiana* epidermal cells. Scale bars: 10 μm.

To determine whether the association between ABA receptors and MAG5 was maintained following perturbation of membrane trafficking, seedlings were treated with BFA, which induces the formation of BFA compartments by disrupting Golgi-TGN/EE (trans-Golgi network/early endosome) trafficking, or WM, which causes the swelling of multivesicular bodies/ prevacuolar compartments (MVBs/PVCs) (Ritzenthaler *et al*., 2002; Tse *et al*., 2004). Following BFA treatment, MAG5 and PYR1 accumulated within the same BFA-induced compartments, which were typically stained by FM4-64 (Fig. **2b**). Likewise, following WM treatment MAG5 remained closely associated with PYR1 in enlarged LysoTracker-positive acidic compartments, consistent with the WM-induced swelling of MVBs/PVCs (Fig. **2b**) (Tse *et al*., 2004; Xu *et al*., 2014). These observations indicate that the spatial association between ABA receptors and MAG5 is preserved following pharmacological perturbation of membrane trafficking. Interestingly, although MAG5 is primarily regarded as an organizer of ER exit sites, it was consistently detected in BFA-induced compartments and WM-induced late endosomal structures together with PYR1, suggesting that its association with PYR1 is maintained beyond steady-state ER exit sites during pharmacological perturbation of membrane trafficking.

To further assess the in *vivo* association between ABA receptors and MAG5, BiFC assays were performed in *N. benthamiana* leaves. Reconstituted YFP fluorescence was detected for both the MAG5-PYR1 and MAG5-PYL4 combinations, whereas no signal was observed in the corresponding negative controls (Figs **2c****, S4**). The BiFC signal exhibited a reticular pattern together with discrete punctate structures that largely overlapped with the ER marker ChFP-KDEL, indicating that the association occurs at ER-associated membrane domains. Although BiFC does not distinguish between direct physical interaction and indirect association within protein complexes or membrane domains, these observations are consistent with a close *in vivo* association between MAG5 and ABA receptors.

The close association between ABA receptors and MAG5 suggested that MAG5 is functionally required for receptor trafficking. To test this hypothesis, we compared the subcellular distribution and trafficking of PYR1 and PYL4 in wild-type and *mag5-3* mutant plants (Takagi *et al*., 2013). Comparison of wild-type and *mag5-3* seedlings revealed an apparent alteration in the steady-state distribution of PYR1 and PYL4, characterized by a more prominent reticular pattern (Fig. **3a**), indicating that disruption of MAG5-dependent ER export alters the normal subcellular organization of ABA receptors.

**Figure 3.**
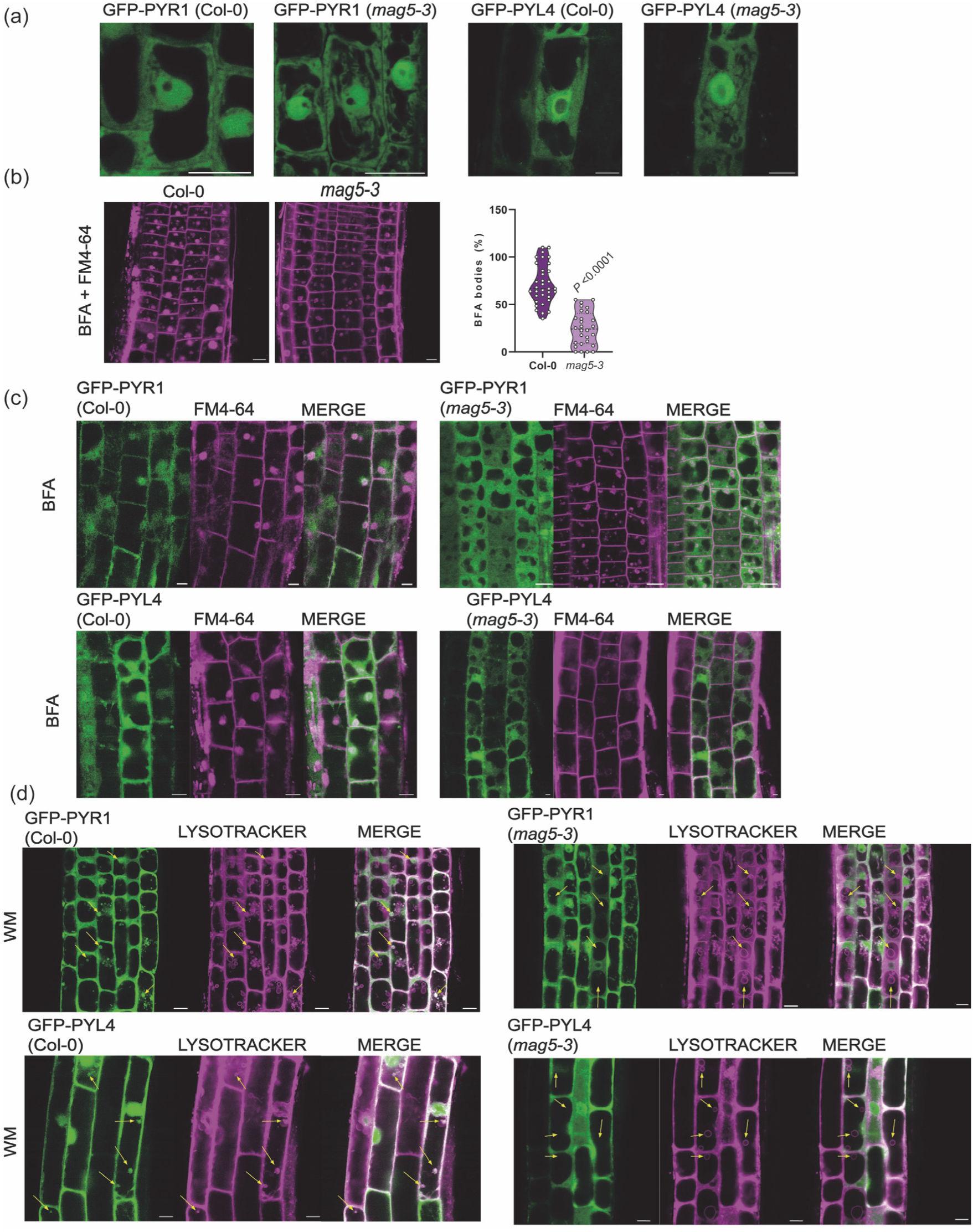
MAG5 is required for the normal trafficking of ABA receptors through the endomembrane system. a) Representative confocal images of root epidermal cells from wild-type (Col-0) and *mag5-3* seedlings expressing *35S::GFP-PYR1* or *35S::GFP–PYL4*. (b) Representative confocal images showing FM4-64-labelled BFA bodies in wild-type (Col-0) and *mag5-3* root epidermal cells following treatment with 50 μM BFA for 1 h and subsequent staining with FM4-64. The graph on the right shows the percentage of cells containing characteristic BFA bodies, defined as well-rounded FM4-64-positive structures larger than 2.5 μm in diameter. Data represent the mean ± SEM (Col-0, n=32 images containing 853 cells from 10 plants; *mag5-3*, n=26 images containing 961 cells from 10 plants). P-values (*P*) denote statistically significant differences (Student’s t-test). (c) Representative confocal images of root epidermal cells expressing *35S::GFP–PYR1* or *35S::GFP–PYL4* in Col-0 or *mag5-3* background following BFA treatment and FM4-64 staining. (d) Representative confocal images of root epidermal cells expressing *35S::GFP–PYR1* or *35S::GFP–PYL4* in Col-0 or *mag5-3* background, following treatment with 33 μM WM for 1 h and staining with LysoTracker™ Deep Red. Arrows indicate representative WM-induced LysoTracker-positive late endosomal compartments. Scale bars: 10 μm

Given the established role of MAG5 in organizing ER exit sites and maintaining ER– Golgi interface integrity (Takagi *et al*., 2013), we next examined whether disruption of MAG5 altered the organization of downstream endomembrane compartments by analysing BFA-induced membrane reorganization. FM4-64 staining revealed the formation of characteristic, well-rounded BFA bodies in wild-type root cells. In contrast, BFA body formation was markedly altered in the *mag5-3* mutant, resulting in fewer, smaller and morphologically aberrant BFA bodies (Fig. **3b**), indicating that disruption of MAG5 alters the organization of BFA-induced endomembrane compartments. In agreement with the altered morphology and reduced formation of BFA bodies, redistribution of GFP–PYR1 and GFP–PYL4 into FM4-64-labelled BFA bodies appeared less pronounced in the *mag5-3* mutant, where receptor fluorescence remained more diffusely distributed throughout the cell (Fig. **3c**).

To determine whether MAG5-dependent ER export influences receptor trafficking towards late endosomal compartments, seedlings were treated with WM. In wild-type plants, PYR1 and PYL4 accumulated within enlarged LysoTracker-positive acidic structures induced by WM (Fig. **3d**), consistent with their delivery to late endosomal MVBs/PVCs (Takáč *et al*., 2012). In contrast, accumulation of PYR1 within these structures was markedly reduced in the *mag5-3* mutant, indicating that disruption of MAG5 impairs the normal trafficking of ABA receptors towards late endosomal compartments, a pathway previously implicated in the vacuolar degradation of ABA receptors (Belda-Palazon *et al*., 2016; Yu *et al*., 2016; García-León *et al*., 2019; Moulinier-Anzola *et al*., 2024; Chen *et al*., 2025). Together, these observations demonstrate that MAG5 is required for the normal intracellular trafficking of ABA receptors through the endomembrane system, leading to their delivery to late endosomal compartments.

### MAG5-dependent trafficking regulates ABA receptor abundance

The altered intracellular trafficking of ABA receptors observed in the *mag5-3* mutant prompted us to examine whether disruption of MAG5 also affects receptor abundance. Immunoblot analysis revealed a significant increase in both PYR1 and PYL4 protein levels in the *mag5-3* mutant compared with wild-type plants under control conditions (Fig. **4a,b**).

**Figure 4.**
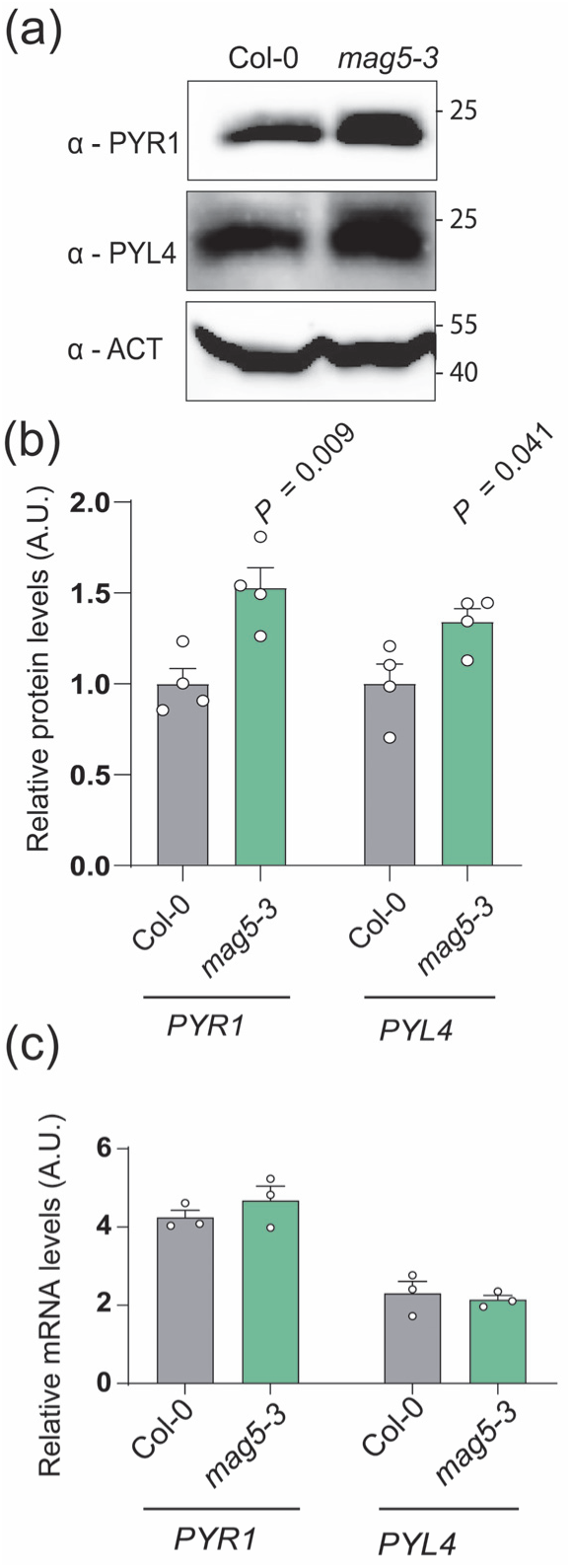
MAG5-dependent trafficking regulates ABA receptor abundance through a post-transcriptional mechanism. (a) Immunoblot analysis of PYR1 and PYL4 protein accumulation in wild-type (Col-0) and *mag5-3* seedlings using the indicated antibodies. ACTIN was used as a loading control. (b) Densitometric quantification of PYR1 and PYL4 protein levels shown in (a). Protein abundance was normalized to ACTIN and referenced to wild-type (Col-0). Data represent the mean ± SEM (n = 4 independent biological replicates). P-values (*P*) denote statistically significant differences (Student’s t-test). (c) Relative transcript levels of PYR1 and PYL4 in wild-type (Col-0) and *mag5-3* seedlings determined by RT-qPCR. Gene expression was normalized to PDF2. Data represent the mean ± SEM (n = 3 independent biological replicates).

To determine whether the increased receptor abundance reflected changes in gene expression, transcript levels of PYR1 and PYL4 were analysed by RT-qPCR. In contrast to the marked increase in protein accumulation, transcript abundance did not differ significantly between wild-type and *mag5-3* plants (Figs **4c**).

Together, these results indicate that MAG5-dependent trafficking regulates ABA receptor abundance through a post-transcriptional mechanism.

### Loss of MAG5 enhances ABA signalling and ABA responses

To investigate the functional consequences of MAG5 loss of function on ABA signalling, we analysed ABA-dependent developmental and physiological responses using well-characterized ABA-hypersensitive (*hab1-1 abi1-2*) or ABA-insensitive (*snrk2.2 snrk2.3*) mutants as controls (Saez *et al*., 2006; Fujii *et al*., 2007). Compared with wild-type plants, the *mag5-3* mutant displayed enhanced sensitivity to exogenous ABA, as evidenced by reduced radicle emergence, cotyledon greening and slightly stronger inhibition of primary root growth (Fig. **5a,b**). Similar ABA hypersensitivity was also observed in two additional independent *mag5* alleles (*mag5-1* and *mag5-2*). By contrast, the closely related *sec16b-1* mutant exhibited a substantially weaker phenotype than *mag5* alleles, consistent with the lower expression of SEC16B relative to MAG5 reported previously (Fig. **S5**) (Takagi *et al*., 2013). Together, these results indicate that enhanced ABA responses are predominantly associated with disruption of MAG5 function.

**Figure 5.**
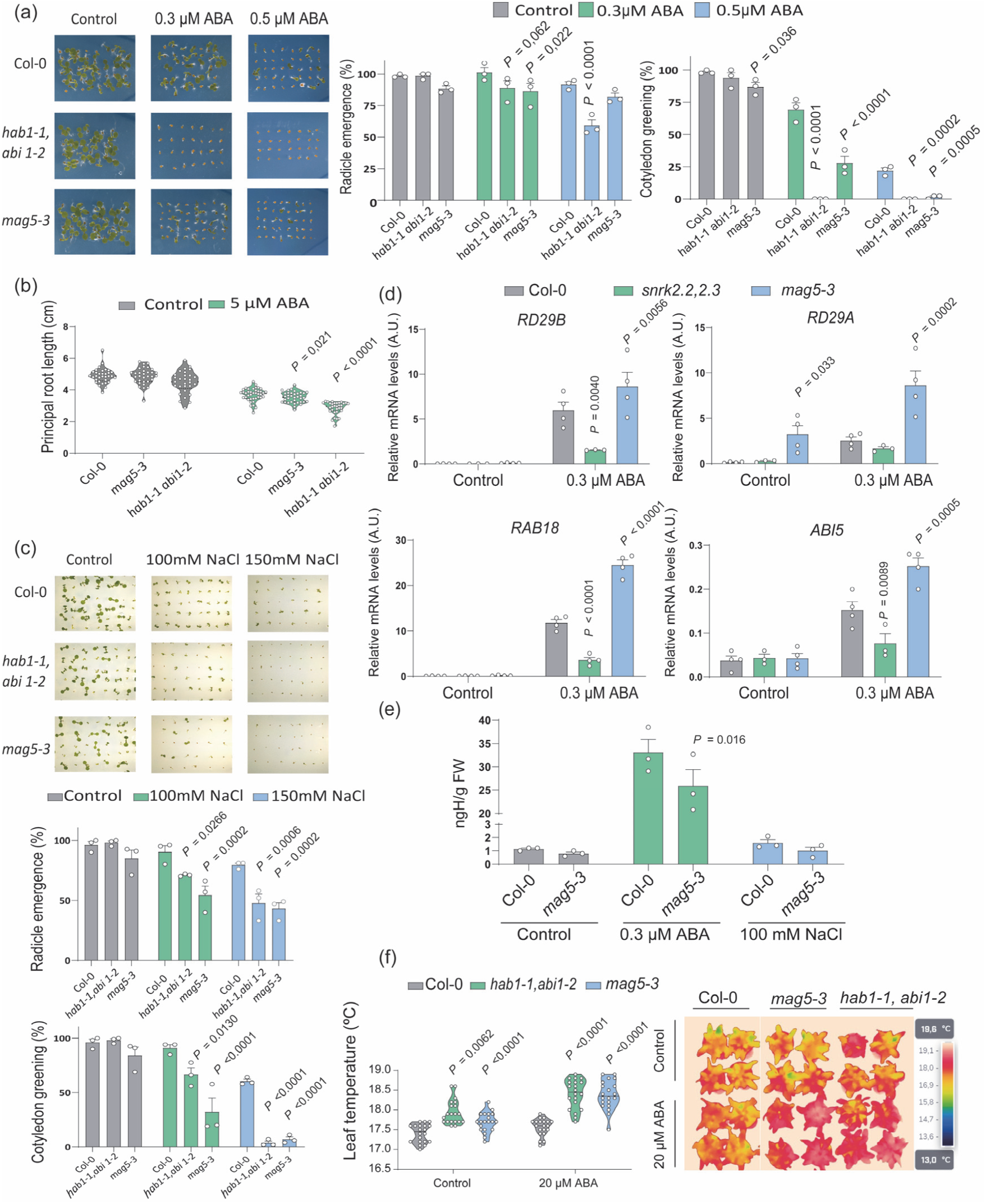
Loss of MAG5 enhances ABA signalling without altering endogenous ABA accumulation. (a) ABA sensitivity during seed germination. Representative images of wild-type (Col-0), the ABA-hypersensitive mutant *hab1-1 abi1-2*, and *mag5-3* seedlings grown on control conditions or media supplemented with 0.3 or 0.5 µM ABA. Quantification of radicle emergence (left) and cotyledon greening (right) of the genotypes is shown. Data represent the mean ± SEM (n = 3 independent biological replicates). P-values (*P*) denote statistically significant differences from wild-type (Col-0) within the same treatment (two-way ANOVA followed by Dunnett’s multiple comparisons test). (b) Primary root growth of wild-type (Col-0), *mag5-3*, and *hab1-1 abi1-2* seedlings grown on control medium or medium supplemented with 5 µM ABA. Data are presented as mean ± SEM (n=29-40 seedlings). P-values (*P*) denote statistically significant differences from wild-type (Col-0) within the same treatment (Student’s t-test). (c) Salt sensitivity during early seedling establishment. Representative images of wild-type (Col-0), *hab1-1 abi1-2*, and *mag5-3* seedlings grown on control medium or supplemented with 100 and 150 mM NaCl. Quantification of radicle emergence (left) and cotyledon greening (right) is shown below. Data represent the mean ± SEM (n = 3 independent biological replicates). P-values (*P*) denote statistically significant differences from wild-type (Col-0) within the same treatment (two-way ANOVA followed by Dunnett’s multiple comparisons test). (d) ABA-responsive gene expression. Relative transcript levels of *RD29B*, *RD29A*, *RAB18* and *ABI5* determined by RT-qPCR in wild-type (Col-0), the ABA-insensitive mutant *snrk2.2 snrk2.3*, and *mag5-3* seedlings grown for 6 d on control medium or supplemented with 0.3 μM ABA. Data represent the mean ± SEM (n = 4 independent biological replicates). P-values (*P*) denote statistically significant differences from wild-type (Col-0) within the same treatment (two-way ANOVA followed by Dunnett’s multiple comparisons test). (e) Endogenous ABA quantification in wild-type (Col-0) and *mag5-3* seedlings under the same treatments indicated in (a) and (c). P-values (*P*) denote statistically significant differences from wild-type (Col-0) within the same treatment (Student’s t-test). (f) Infrared thermography analysis of rosette leaf temperature in wild-type (Col-0), *hab1-1 abi1-2*, and *mag5-3* plants following control or ABA treatment (20 µM). Representative thermal images are shown together with the corresponding quantification of leaf temperature. Same infrared thermography image was cropped for showing wild-type (Col-0) and *mag5-3* contiguously. Violin box plots representing the leaf measurements of 18 plants. P-values (*P*) denote statistically significant differences from wild-type (Col-0) within the same treatment (two-way ANOVA followed by Dunnett’s multiple comparisons test).

To determine whether this enhanced ABA responsiveness also affected stress adaptation, we examined early growth under salt stress. In agreement with their ABA-hypersensitive phenotype, *mag5-3* seedlings displayed a stronger inhibition of radicle emergence and cotyledon greening in the presence of 100 and 150 mM NaCl, closely resembling the ABA-hypersensitive *hab1-1 abi1-2* mutant, whereas wild-type seedlings were considerably less affected (Fig. **5c**). These results indicate that disruption of MAG5 enhances physiological responses to salt stress, in line with increased ABA signalling.

To determine whether the enhanced ABA sensitivity of *mag5-3* was accompanied by increased activation of ABA signalling, we analysed the expression of the ABA-responsive marker genes *RD29A*, *RD29B*, *ABI5* and *RAB18* (Fujii *et al*., 2007). Under mock conditions, transcript levels were similarly low in wild-type, *mag5-3* and the ABA-insensitive *snrk2.2 snrk2.3* mutant (Fig. **5d**). Following ABA treatment, *RD29A*, *RD29B*, *ABI5* and *RAB18* were strongly induced in wild-type seedlings, whereas induction was almost completely abolished in *snrk2.2 snrk2.3*, confirming the effectiveness of the treatment. By contrast, ABA-induced expression of both marker genes was significantly higher in *mag5-3* than in wild-type plants (Fig. **5d**), demonstrating that disruption of MAG5 enhances ABA signalling at the transcriptional level.

To exclude the possibility that the enhanced ABA sensitivity of *mag5-3* resulted from altered endogenous ABA accumulation, we quantified ABA levels under control, exogenous ABA treatment and salt stress conditions. Endogenous ABA concentrations were comparable between wild-type and *mag5-3* seedlings under control conditions and following salt treatment. Upon treatment with 0.3 µM ABA, however, *mag5-3* seedlings accumulated significantly less ABA than wild-type seedlings (Fig. **5e**). Thus, the enhanced ABA sensitivity of *mag5-3* cannot be explained by increased ABA accumulation. Instead, these results support the conclusion that disruption of MAG5 enhances ABA responsiveness at the level of signal perception or transduction rather than hormone accumulation.

Because ABA promotes stomatal closure, thereby reducing transpirational cooling and increasing leaf temperature, we next examined whether enhanced ABA signalling in *mag5-3* was associated with altered stomatal responses using infrared thermography (Jones & Leinonen, 2003; Wang *et al*., 2004; Munemasa *et al*., 2015; Vaidya *et al*., 2019). Infrared thermography revealed that *mag5-3* leaves exhibited significantly higher leaf temperatures than wild-type plants under both mock and ABA-treated conditions (Fig. **5f**). Notably, *mag5-3* displayed elevated leaf temperatures even in the absence of exogenous ABA, suggesting enhanced responsiveness to endogenous ABA. Following ABA treatment, leaf temperature increased in all genotypes but was significantly higher in *mag5-3* than in wild-type plants, approaching the response observed in the ABA-hypersensitive control *hab1-1 abi1-2*. To directly assess whether this thermographic phenotype was associated with altered stomatal behaviour, we measured stomatal aperture in seedling cotyledons. Consistent with the thermographic phenotype, *mag5-3* seedlings displayed a significantly reduced stomatal aperture compared with Col-0 (Fig. **S6**). Together, these results support enhanced stomatal closure in the absence of MAG5 and provide physiological evidence that altered early secretory trafficking affects not only growth-related ABA responses but also stomatal regulation.

## Discussion

ABA receptor homeostasis depends on the coordinated regulation of intracellular trafficking and protein turnover. Over the past decade, considerable progress has identified multiple ubiquitin ligases and endosomal sorting components that control receptor abundance through proteasomal or vacuolar degradation (Coego *et al*., 2021; Qin *et al*., 2026). In contrast, whether early secretory trafficking contributes to receptor homeostasis has remained largely unexplored. Our findings identify MAG5-dependent ER export as a regulatory checkpoint contributing to ABA receptor homeostasis by controlling receptor trafficking at the earliest stages of the secretory pathway (Fig. **6**).

**Figure 6.**
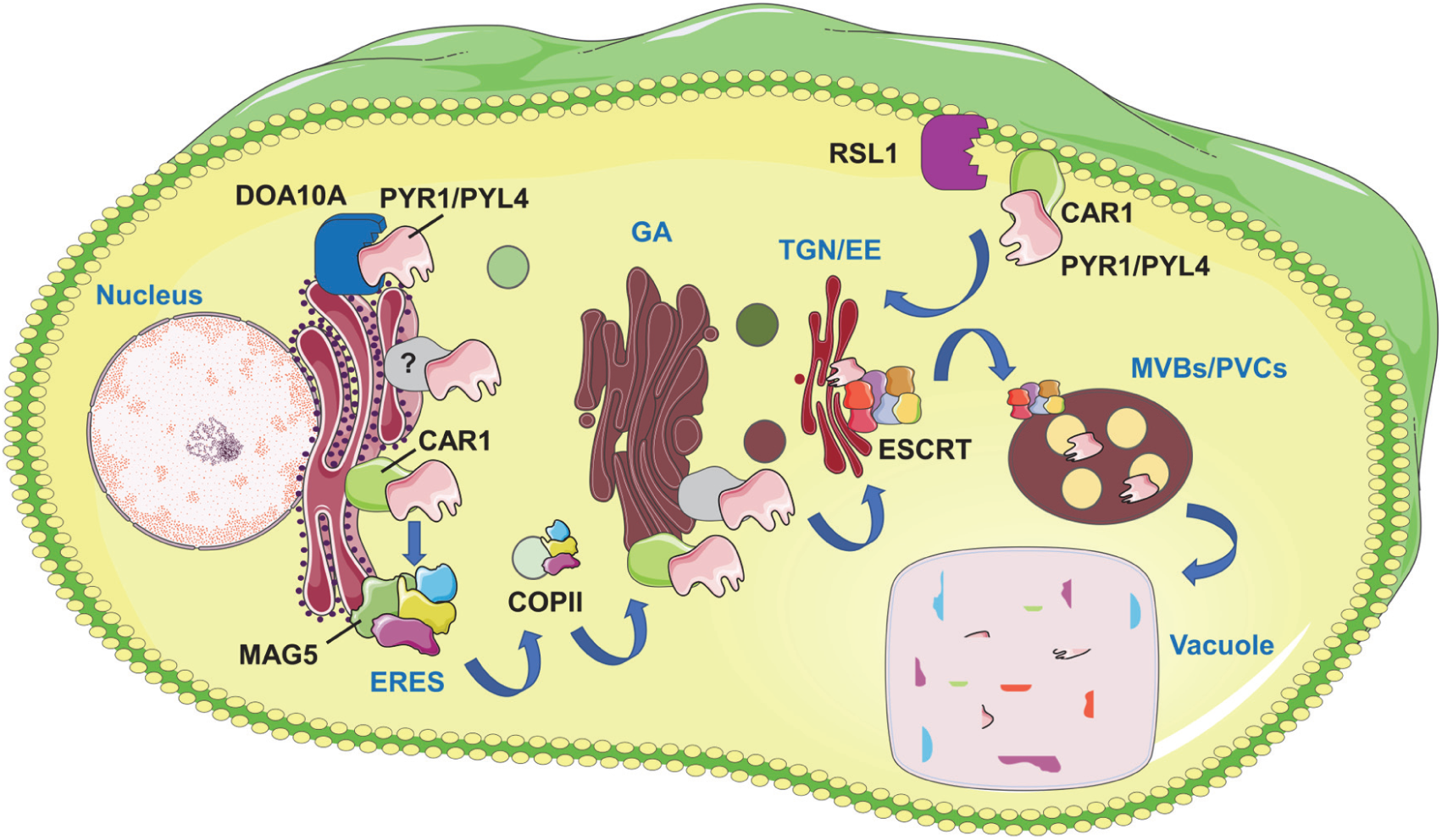
Proposed model for the role of MAG5-dependent ER export in the establishment of ABA receptor homeostasis. ABA receptors are proposed to maintain a constitutive pool associated with the ER, where receptor abundance and trafficking competence may already be regulated through interactions with quality-control and trafficking factors, including the ER-associated E3 ubiquitin ligase DOA10A and CAR proteins. Receptors that successfully exit the ER through MAG5- and COPII-dependent ER exit sites (ERES) are subsequently delivered to the Golgi apparatus (GA) and the trans-Golgi network/early endosome (TGN/EE). During their progression through the secretory pathway, receptors may associate with additional regulatory factors before being sorted either to the plasma membrane, where RSL1 promotes receptor ubiquitination and endocytosis, or into the endosomal pathway, leading to multivesicular bodies/prevacuolar compartments (MVBs/PVCs) and ESCRT-dependent vacuolar degradation. Disruption of MAG5-dependent ER export is proposed to alter receptor distribution throughout the endomembrane system while limiting progression towards endosomal sorting and vacuolar degradation. Such redistribution increases the proportion of receptors available for ABA perception and signalling, thereby enhancing ABA responsiveness without affecting endogenous ABA accumulation.

Based on our findings and previous studies (Coego *et al*., 2021; Li *et al*., 2021; Cui *et al*., 2023; Liu *et al*., 2025), we propose a model in which a constitutive pool of ABA receptors is maintained at the ER, where receptor abundance and trafficking competence may already be regulated through interactions with quality-control and trafficking factors. Receptors that successfully exit the ER through MAG5- and COPII-dependent ER exit sites are subsequently delivered to the Golgi apparatus (GA) and the TGN. Along this secretory itinerary, receptors may acquire additional regulatory partners, including CAR proteins, before being sorted either to the plasma membrane or into the endosomal pathway, ultimately reaching MVBs/PVCs and the vacuole for degradation through ESCRT-dependent trafficking. Disruption of MAG5 is predicted to alter the normal distribution of ABA receptors across the endomembrane system by reducing the efficient delivery of ABA receptors to downstream endomembrane compartments while simultaneously limiting their progression towards endosomal sorting and vacuolar degradation. Such redistribution would alter the balance between receptor pools available for ABA perception and signalling but also, those committed to degradation, providing a plausible explanation for the increased accumulation of PYR1 and PYL4 observed in *mag5-3* mutants despite impaired trafficking through the secretory pathway. Together, this model provides a conceptual framework linking early secretory trafficking with the establishment of ABA receptor homeostasis across the endomembrane system (Fig. **6**).

Consistent with this model, disruption of another core COPII component, SAR1A, also results in enhanced ABA sensitivity, suggesting that COPII-dependent ER export represents a broader regulatory layer controlling ABA signalling rather than a mechanism unique to MAG5. Our findings are particularly intriguing in light of this work, demonstrating that ABA actively remodels the early secretory pathway by inducing SAR1A expression and promoting the formation of specialized COPII vesicles required for the trafficking of stress-associated membrane proteins (Li *et al*., 2021). Together with these observations, our data support a reciprocal regulatory relationship between ABA signalling and the early secretory pathway. Whereas ABA signalling enhances COPII function through SAR1A-dependent remodelling of ER export, our results indicate that efficient COPII-mediated trafficking is required to establish the abundance and intracellular distribution of ABA receptors. Moreover, our pharmacological analyses indicate that the consequences of MAG5 disruption extend well beyond ER exit sites. In addition to the reduced accumulation of PYR1 and PYL4 in WM-induced late endosomal compartments, the altered morphology and reduced formation of BFA bodies in *mag5-3* are in line with the established role of MAG5 in maintaining the organization of ER exit sites and efficient ER export (Takagi et al., 2013). Rather than reflecting a direct defect in endosomal trafficking, these observations are more consistent with a global perturbation of membrane flux across the endomembrane system. Previous studies of the ER–Golgi tethering proteins MAG2 and MAG4, together with the ER exit site organizer MAG5, demonstrated that disruption of distinct trafficking steps at the ER–Golgi interface leads to a common phenotype characterized by defective cargo export, accumulation of secretory cargo within ER-derived MAG bodies, and impaired delivery of vacuolar proteins to protein storage vacuoles (Li *et al*., 2006; Takahashi *et al*., 2010; Takagi *et al*., 2013). These studies further suggested that defects in early secretory trafficking disrupt the dynamic balance between anterograde and retrograde membrane transport. Within this framework, impaired COPII-dependent ER export would reduce the continuous supply of membrane and cargo to downstream compartments, thereby progressively altering the organization and trafficking capacity of the Golgi and TGN/EE system. Consequently, ABA receptors would be delivered less efficiently to late endosomes and vacuoles, reducing their progression through degradative pathways while simultaneously favoring their accumulation in earlier secretory compartments.

This interpretation is in agreement with the emerging view that the endomembrane system operates as an integrated trafficking network, in which perturbations arising at early secretory steps propagate throughout the endomembrane system and ultimately influence downstream trafficking events. Our observations fit well within this conceptual framework, as defects in MAG5-dependent ER export are accompanied by altered organization of BFA-sensitive compartments and reduced accumulation of ABA receptors in WM-induced late endosomal structures. In line with this idea, recent studies indicate that COPII components participate in membrane remodelling events extending beyond conventional ER-to-Golgi transport, revealing unexpected functional connections between ER exit sites and downstream degradative membrane systems (Liao *et al*., 2024; Chung *et al*., 2024). Although our observations do not demonstrate a direct role for MAG5 in late endosomal trafficking, they support the emerging view that early secretory trafficking and endosomal sorting are functionally integrated processes rather than independent transport pathways. This bidirectional interplay reinforces the concept that the early secretory pathway is not merely a constitutive transport route but a dynamic signalling hub that both responds to and shapes hormone signalling (Aniento *et al*., 2022).

An emerging theme in ABA signalling is that receptor homeostasis is coordinated in a compartment-specific manner rather than through a single degradation pathway (Coego *et al*., 2021; Qin *et al*., 2026). Different E3 ubiquitin ligases regulate distinct subcellular pools of ABA receptors, collectively establishing spatial control over receptor abundance and turnover. For instance, the plasma membrane-associated RSL1 promotes ubiquitination and endocytosis of PYR1 and PYL4, leading to their subsequent sorting through the ESCRT pathway for vacuolar degradation, whereas the related RFA family members regulate receptor turnover in the cytosol and nucleus through proteasomal degradation. Likewise, the CRL4-DDA1 complex targets nuclear ABA receptors, whereas the ER-associated E3 ligase DOA10A promotes receptor mono-ubiquitination and relocalization towards membrane compartments (Liu *et al*., 2025). Our findings extend this compartmentalized view of receptor regulation by placing the ER–Golgi interface upstream of the previously described compartment-specific regulatory mechanisms.

Importantly, the physiological consequences of ABA receptor trafficking extend beyond the control of receptor abundance itself. Previous work on FYVE1/FREE1, FYVE4, VPS23A, ALIX and TOL proteins has shown that perturbation of receptor endocytosis, endosomal sorting and vacuolar degradation alters ABA receptor homeostasis and hormone responsiveness, affecting both developmental and physiological ABA responses, including stomatal regulation (Belda-Palazon *et al*., 2016; Yu *et al*., 2016; García-León *et al*., 2019; Moulinier-Anzola *et al*., 2024; Chen *et al*., 2025). This is particularly relevant because ABA signalling controls not only transcriptional and growth responses but also rapid membrane-delimited processes required for adaptation to water stress. The recruitment of specific ABA receptors to the plasma membrane by CAR proteins and the DOA10A-dependent regulation of receptor trafficking towards membrane compartments further highlight the importance of receptor positioning for ABA signalling (Diaz *et al*., 2016; Liu *et al*., 2025). In this context, our finding that disruption of MAG5 alters ABA receptor distribution and affects both growth-related ABA responses and stomatal regulation extends the functional relevance of receptor trafficking to the early secretory pathway. Although our data do not directly establish how MAG5-dependent ER export affects receptor abundance at the plasma membrane, they support a model in which perturbations at the ER–Golgi interface alter receptor distribution and ultimately influence distinct ABA signalling outputs.

A major implication of our findings is the identification of a constitutive ER-associated pool of ABA receptors, indicating that the ER functions as an active regulatory compartment rather than merely a transient station of the secretory pathway. In this context, receptor export is likely coordinated with additional processes, including post-translational modifications such as DOA10A-mediated mono-ubiquitination and the association with CAR proteins, which together may determine receptor trafficking competence and progression through COPII-dependent ER export. Defining how these mechanisms are integrated with COPII-mediated trafficking will be essential for understanding how receptor homeostasis is dynamically regulated.

Collectively, our work expands current models of ABA signalling by demonstrating that receptor trafficking is an integral determinant of hormone responsiveness, revealing that the ER–Golgi interface contributes directly to the establishment of ABA receptor homeostasis.

## Acknowledgements

We thank Ikuko Hara-Nishimura for the *pMAG5::MAG5-GFP*, *mag5-1*, *mag5-2*, *mag5-3* and *sec16b-1 A. thaliana* transgenic lines. The authors acknowledge the assistance of the IBMCP Scientific and Technical Services, including the Advanced Microscopy Unit, the Plant Growth Facility and the Hormone Quantification Facility, for their support during the course of this work. We gratefully acknowledge the financial support that made this study possible. This research was funded by the Spanish Ministry of Science, Innovation and Universities (MCIU), grant numbers PID2022-142412NB-I00 (awarded to B.B.-P. and A.F.), PID2020-113847GB-I00 (awarded de F.A and M.J.M.), and CNS2024-154566 (awarded to B.B.-P.), and by the CIDEGENT program of excellence of GVA, grant number CIDEXG/2022/27 (awarded to B.B.-P.). This work was also supported by grant PID2023-147322OB-I00, funded by MCIU/AEI/10.13039/501100011033 and the ERDF (“A way of making Europe”) (awarded to P.L.R.). B.K. and J.M.-A. acknowledge funding from the Austrian Science Fund (FWF) through grants PAT1121425 and P34748. J.A.B.-T. and A.V.-F. were respectively the beneficiary of PhD contracts CIDEXG/2022/27-01 and CIDEXG/2022/27-02, funded by GVA. We also acknowledge financial support from the institutional grant CEX2025-001626-S (Center of Excellence Severo Ochoa) funded by MCIU/AEI/10.13039/501100011033.

## Competing interests

None declared.

## Author contributions

J.A.B.-T., A.V.-F., C.C.-R. M.B., R.M. and B.B.-P. conducted experiments. F.A., M.J.M., J.M.-A., B.K., P.L.R. provided tools and expertise on vesicle trafficking and ABA signalling, actively supporting the conceptual work. B.B.-P. and A.F. conceived and designed the project. B.B.-P. directed and supervised all research activities and prepared figures. B.B.-P. wrote the manuscript. F.A., M.J.M., J.M.-A., B.K., P.L.R and A.F. reviewed the manuscript. All authors have read and agreed to the published version of the manuscript.

## Data availability

All data supporting the findings of this study are available within the article and its Supporting Information. Additional data and materials related to this study are available from the corresponding author upon reasonable request.

## Notes

### Competing Interest Statement

The authors have declared no competing interest.

## References

Aniento F, de Medina Hernández VS, Dagdas Y, Rojas-Pierce M, Russinova E. 2022. Molecular mechanisms of endomembrane trafficking in plants. The Plant Cell 34: 146–173.

Belda-Palazón B, Rodriguez PL. 2020. Degradation of Abscisic Acid Receptors Through the Endosomal Pathway. *Methods in molecular biology (Clifton*, N.J.) 2177: 35– 48.

Belda-Palazón B, Rodriguez PL. 2022. Microscopic Imaging of Endosomal Trafficking of ABA Receptors. *Methods in molecular biology (Clifton*, N.J.) 2462: 59–69.

Belda-Palazon B, Rodriguez L, Fernandez MA, Castillo MC, Anderson EM, Gao C, Gonzalez-Guzman M, Peirats-Llobet M, Zhao Q, De Winne N, et al. 2016. FYVE1/FREE1 Interacts with the PYL4 ABA Receptor and Mediates Its Delivery to the Vacuolar Degradation Pathway. The Plant cell 28: 2291–2311.

Belda-Palazón B, Ruiz L, Martí E, Tárraga S, Tiburcio AF, Culiáñez F, Farràs R, Carrasco P, Ferrando A. 2012. Aminopropyltransferases Involved in Polyamine Biosynthesis Localize Preferentially in the Nucleus of Plant Cells. PLoS ONE 7.

Bernat-Silvestre C, Ma Y, Johnson K, Ferrando A, Aniento F, Marcote MJ. 2022. Characterization of Arabidopsis Post-Glycosylphosphatidylinositol Attachment to Proteins Phospholipase 3 Like Genes. Frontiers in Plant Science 13: 817915.

Brandizzi F. 2018. Transport from the endoplasmic reticulum to the Golgi in plants: Where are we now? Seminars in Cell & Developmental Biology 80: 94–105.

Brandizzi F, Barlowe C. 2013. Organization of the ER-Golgi interface for membrane traffic control. Nature Reviews Molecular Cell Biology 14: 382–392.

Bueso E, Rodriguez L, Lorenzo-Orts L, Gonzalez-Guzman M, Sayas E, Muñoz-Bertomeu J, Ibañez C, Serrano R, Rodriguez PL. 2014. The single-subunit RING-type E3 ubiquitin ligase RSL1 targets PYL4 and PYR1 ABA receptors in plasma membrane to modulate abscisic acid signaling. Plant Journal 80: 1057–1071.

Chen J, Liu C, Yang Z, Zhuo D, Li H, Zhou J, Yang C, Shen W, Peng C, Gao C. 2025. The endosomal sorting complex protein FYVE4 affects abscisic acid signaling by regulating the stability of abscisic acid receptors. Journal of Experimental Botany 76: 6185–6198.

Chung T, Choi YE, Song K, Jung H. 2024. How coat proteins shape autophagy in plant cells. Plant Physiology 197.

Clough SJ, Bent AF. 1998. Floral dip: A simplified method for Agrobacterium-mediated transformation of Arabidopsis thaliana. Plant Journal 16: 735–743.

Coego A, Julian J, Lozano-Juste J, Pizzio GA, Alrefaei AF, Rodriguez PL. 2021. Ubiquitylation of ABA Receptors and Protein Phosphatase 2C Coreceptors to Modulate ABA Signaling and Stress Response. International Journal of Molecular Sciences 2021, *Vol. 22, Page 7103 22: 7103*.

Connerly PL, Esaki M, Montegna EA, Strongin DE, Levi S, Soderholm J, Glick BS. 2005. Sec16 is a Determinant of Transitional ER Organization. Current Biology 15: 1439–1447.

Cui M, Gupta SK, Bauer P. 2023. Role of the plant-specific calcium-binding C2-DOMAIN ABSCISIC ACID-RELATED (CAR) protein family in environmental signaling. European Journal of Cell Biology 102: 151322.

Cutler SR, Rodriguez PL, Finkelstein RR, Abrams SR. 2010. Abscisic Acid: Emergence of a Core Signaling Network. 10.1146/annurev-arplant-042809-112122 61: 651–679.

Diaz M, Sanchez-Barrena MJ, Gonzalez-Rubio JM, Rodriguez L, Fernandez D, Antoni R, Yunta C, Belda-Palazon B, Gonzalez-Guzman M, Peirats-Llobet M, et al. 2016. Calcium-dependent oligomerization of CAR proteins at cell membrane modulates ABA signaling. Proceedings of the National Academy of Sciences of the United States of America 113: E396–E405.

Fernandez MA, Belda-Palazon B, Julian J, Coego A, Lozano-Juste J, Iñigo S, Rodriguez L, Bueso E, Goossens A, Rodriguez PL. 2020. RBR-Type E3 Ligases and the Ubiquitin-Conjugating Enzyme UBC26 Regulate Abscisic Acid Receptor Levels and Signaling. Plant physiology 182: 1723–1742.

Fujii H, Verslues PE, Zhu J-K. 2007. Identification of Two Protein Kinases Required for Abscisic Acid Regulation of Seed Germination, Root Growth, and Gene Expression in Arabidopsis. The Plant Cell 19: 485–494.

García-León M, Cuyas L, Abd El-Moneim D, Rodriguez L, Belda-Palazón B, Sanchez-Quant E, Fernández O, Roux B, María Zamarreño Á, María García-Mina J, et al. 2019. Arabidopsis ALIX Regulates Stomatal Aperture and Turnover of Abscisic Acid Receptors. The Plant cell 31: 2411–2429.

Geiger D, Scherzer S, Mumm P, Stange A, Marten I, Bauer H, Ache P, Matschi S, Liese A, Al-Rasheid KAS, et al. 2009. Activity of guard cell anion channel SLAC1 is controlled by drought-stress signaling kinase-phosphatase pair. Proceedings of the National Academy of Sciences of the United States of America 106: 21425–21430.

Imes D, Mumm P, Böhm J, Al-Rasheid KAS, Marten I, Geiger D, Hedrich R. 2013. Open stomata 1 (OST1) kinase controls R-type anion channel QUAC1 in Arabidopsis guard cells. Plant Journal 74: 372–382.

Irigoyen ML, Iniesto E, Rodriguez L, Puga MI, Yanagawa Y, Pick E, Strickland E, Paz-Ares J, Wei N, De Jaeger G, et al. 2014. Targeted Degradation of Abscisic Acid Receptors Is Mediated by the Ubiquitin Ligase Substrate Adaptor DDA1 in Arabidopsis. The Plant Cell 26: 712–728.

Jones HG, Leinonen I. 2003. Thermal Imaging for the Study of Plant Water Relations. Journal of Agricultural Meteorology 59: 205–217.

Kollist H, Nuhkat M, Roelfsema MRG. 2014. Closing gaps: Linking elements that control stomatal movement. New Phytologist 203: 44–62.

Li L, Shimada T, Takahashi H, Ueda H, Fukao Y, Kondo M, Nishimura M, Hara-Nishimura I. 2006. MAIGO2 Is Involved in Exit of Seed Storage Proteins from the Endoplasmic Reticulum in Arabidopsis thaliana. The Plant Cell 18: 3535–3547.

Li B, Zeng Y, Cao W, Zhang W, Cheng L, Yin H, Wu Q, Wang X, Huang Y, Lau WCY, et al. 2021. A distinct giant coat protein complex II vesicle population in Arabidopsis thaliana. Nature Plants 2021 7:10 7: 1335–1346.

Liao YC, Pang S, Li WP, Shtengel G, Choi H, Schaefer K, Xu CS, Lippincott-Schwartz J. 2024. COPII with ALG2 and ESCRTs control lysosome-dependent microautophagy of ER exit sites. Developmental Cell 59: 1410–1424.e4.

Liu C, Li Q, Shen Z, Xia R, Chen Q, Li X, Ding Y, Yang S, Serino G, Xie Q, et al. 2025. The Arabidopsis E3 ubiquitin ligase DOA10A promotes localization of abscisic acid (ABA) receptors to the membrane through mono-ubiquitination in ABA signaling. New Phytologist 245: 169–182.

Moulinier-Anzola J, Schwihla M, Lugsteiner R, Leibrock N, Feraru MI, Tkachenko I, Luschnig C, Arcalis E, Feraru E, Lozano-Juste J, et al. 2024. Modulation of abscisic acid signaling via endosomal TOL proteins. New Phytologist 243: 1065–1081.

Munemasa S, Hauser F, Park J, Waadt R, Brandt B, Schroeder JI. 2015. Mechanisms of abscisic acid-mediated control of stomatal aperture. Current Opinion in Plant Biology 28: 154–162.

Nelson BK, Cai X, Nebenführ A. 2007. A multicolored set of in vivo organelle markers for co-localization studies in Arabidopsis and other plants. The Plant Journal 51: 1126–1136.

Paez Valencia J, Goodman K, Otegui MS. 2016. Endocytosis and Endosomal Trafficking in Plants. Annual review of plant biology 67: 309–335.

Qin S, Qin Q, Hou S. 2026. ABA receptors: function and post-translational modifications in plants. Journal of Plant Physiology 317: 154692.

Ritzenthaler C, Nebenführ A, Movafeghi A, Stussi-Garaud C, Behnia L, Pimpl P, Staehelin LA, Robinson DG. 2002. Reevaluation of the Effects of Brefeldin A on Plant Cells Using Tobacco Bright Yellow 2 Cells Expressing Golgi-Targeted Green Fluorescent Protein and COPI Antisera. The Plant Cell 14: 237–261.

Rodriguez L, Gonzalez-Guzman M, Diaz M, Rodrigues A, Izquierdo-Garcia AC, Peirats-Llobet M, Fernandez MA, Antoni R, Fernandez D, Marquez JA, et al. 2015. C2-Domain Abscisic Acid-Related Proteins Mediate the Interaction of PYR/PYL/RCAR Abscisic Acid Receptors with the Plasma Membrane and Regulate Abscisic Acid Sensitivity in Arabidopsis. The Plant Cell 26: 4802–4820.

Rodriguez PL, Lozano-Juste J, Albert A. 2019. PYR/PYL/RCAR ABA receptors. Advances in Botanical Research 92: 51–82.

Saez A, Robert N, Maktabi MH, Schroeder JI, Serrano R, Rodriguez PL. 2006. Enhancement of Abscisic Acid Sensitivity and Reduction of Water Consumption in Arabidopsis by Combined Inactivation of the Protein Phosphatases Type 2C ABI1 and HAB1. Plant Physiology 141: 1389–1399.

Schindelin J, Arganda-Carreras I, Frise E, Kaynig V, Longair M, Pietzsch T, Preibisch S, Rueden C, Saalfeld S, Schmid B, et al. 2012. Fiji: an open-source platform for biological-image analysis. Nature Methods 2012 9:7 9: 676–682.

Serra-Soriano M, Pallás V, Navarro JA. 2014. A model for transport of a viral membrane protein through the early secretory pathway: Minimal sequence and endoplasmic reticulum lateral mobility requirements. Plant Journal 77: 863–879.

Takáč T, Pechan T, Šamajová O, Ovečka M, Richter H, Eck C, Niehaus K, Šamaj J. 2012. Wortmannin Treatment Induces Changes in Arabidopsis Root Proteome and Post-Golgi Compartments. Journal of Proteome Research 11: 3127–3142.

Takagi J, Renna L, Takahashi H, Koumoto Y, Tamura K, Stefano G, Fukao Y, Kondo M, Nishimura M, Shimada T, et al. 2013. MAIGO5 Functions in Protein Export from Golgi-Associated Endoplasmic Reticulum Exit Sites in Arabidopsis. The Plant Cell 25: 4658–4675.

Takahashi H, Tamura K, Takagi J, Koumoto Y, Hara-Nishimura I, Shimada T. 2010. MAG4/Atp115 is a Golgi-Localized Tethering Factor that Mediates Efficient Anterograde Transport in Arabidopsis. Plant and Cell Physiology 51: 1777–1787.

Tse YC, Mo B, Hillmer S, Zhao M, Lo SW, Robinson DG, Jiang L. 2004. Identification of Multivesicular Bodies as Prevacuolar Compartments in Nicotiana tabacum BY-2 Cells. The Plant Cell 16: 672–693.

Vaidya AS, Helander JDM, Peterson FC, Elzinga D, Dejonghe W, Kaundal A, Park SY, Xing Z, Mega R, Takeuchi J, et al. 2019. Dynamic control of plant water use using designed ABA receptor agonists. Science 366.

Wang Y, Holroyd G, Hetherington AM, Ng CKY. 2004. Seeing ‘cool’ and ‘hot’— infrared thermography as a tool for non-invasive, high-throughput screening of Arabidopsis guard cell signalling mutants. Journal of Experimental Botany 55: 1187– 1193.

Wang X, Xu M, Gao C, Zeng Y, Cui Y, Shen W, Jiang L. 2020. The roles of endomembrane trafficking in plant abiotic stress responses. Journal of Integrative Plant Biology 62: 55–69.

Watson P, Townley AK, Koka P, Palmer KJ, Stephens DJ. 2006. Sec16 defines endoplasmic reticulum exit sites and is required for secretory cargo export in mammalian cells. Traffic 7: 1678–1687.

Xu M, Liu K, Swaroop M, Sun W, Dehdashti SJ, McKew JC, Zheng W. 2014. A Phenotypic Compound Screening Assay for Lysosomal Storage Diseases. SLAS Discovery 19: 168–175.

Yoshida T, Christmann A, Yamaguchi-Shinozaki K, Grill E, Fernie AR. 2019. Revisiting the Basal Role of ABA – Roles Outside of Stress. Trends in Plant Science 24: 625–635.

Yu F, Lou L, Tian M, Li Q, Ding Y, Cao X, Wu Y, Belda-Palazon B, Rodriguez PL, Yang S, et al. 2016. ESCRT-I Component VPS23A Affects ABA Signaling by Recognizing ABA Receptors for Endosomal Degradation. Molecular Plant 9: 1570– 1582.

