## Supplemental info for "SEC16A/MAG5-mediated ER export shapes ABA receptor homeostasis"

The following Supporting Information is available for this article:

**Video/Movie S1.** Time-lapse confocal imaging *of N. benthamiana* epidermal cells transiently co-expressing GFP-PYR1 (green) and the ER marker ChFP-KDEL (magenta). Fluorescence emitted by chlorophyll is displayed in blue.

**Video/Movie S2.** Time-lapse confocal imaging *of N. benthamiana* epidermal cells transiently co-expressing CAR1-YFP$^{N}$ with YFP$^{C}$-PYR1 (green) and RFP-MAG5 (magenta).

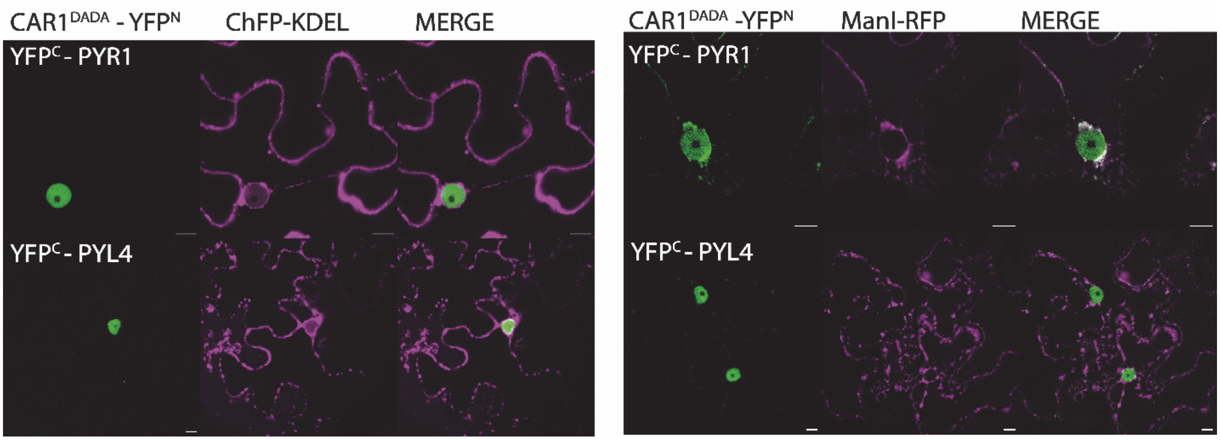


**Figure S1. Disruption of the membrane-binding capacity of CAR1 abolishes the ER- and Golgi-associated BiFC pattern.** Representative BiFC images following transient co-expression in *N. benthamiana* epidermal cells of the membrane-binding-defective mutant CAR1$^{DADA}$-YFP$^{N}$ with YFP$^{C}$-PYR1 or YFP$^{C}$-PYL4 together with ChFP-KDEL (a) or ManI-RFP (b). Scale bars, 10 µm.

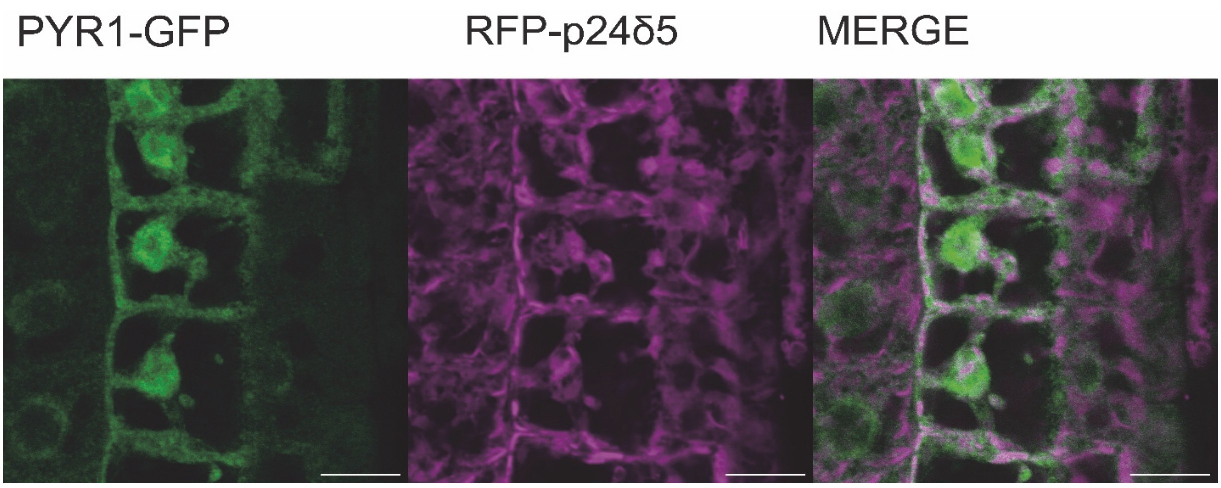


**Figure S2.** Additional confocal images of root epidermal cells from transgenic *A. thaliana* seedlings expressing *pPYR1::PYR1–GFP* together with the ER marker RFP–p24δ5. These images provide additional examples of the co-localization shown in Fig. **1e**. Scale bars, 10 µm.

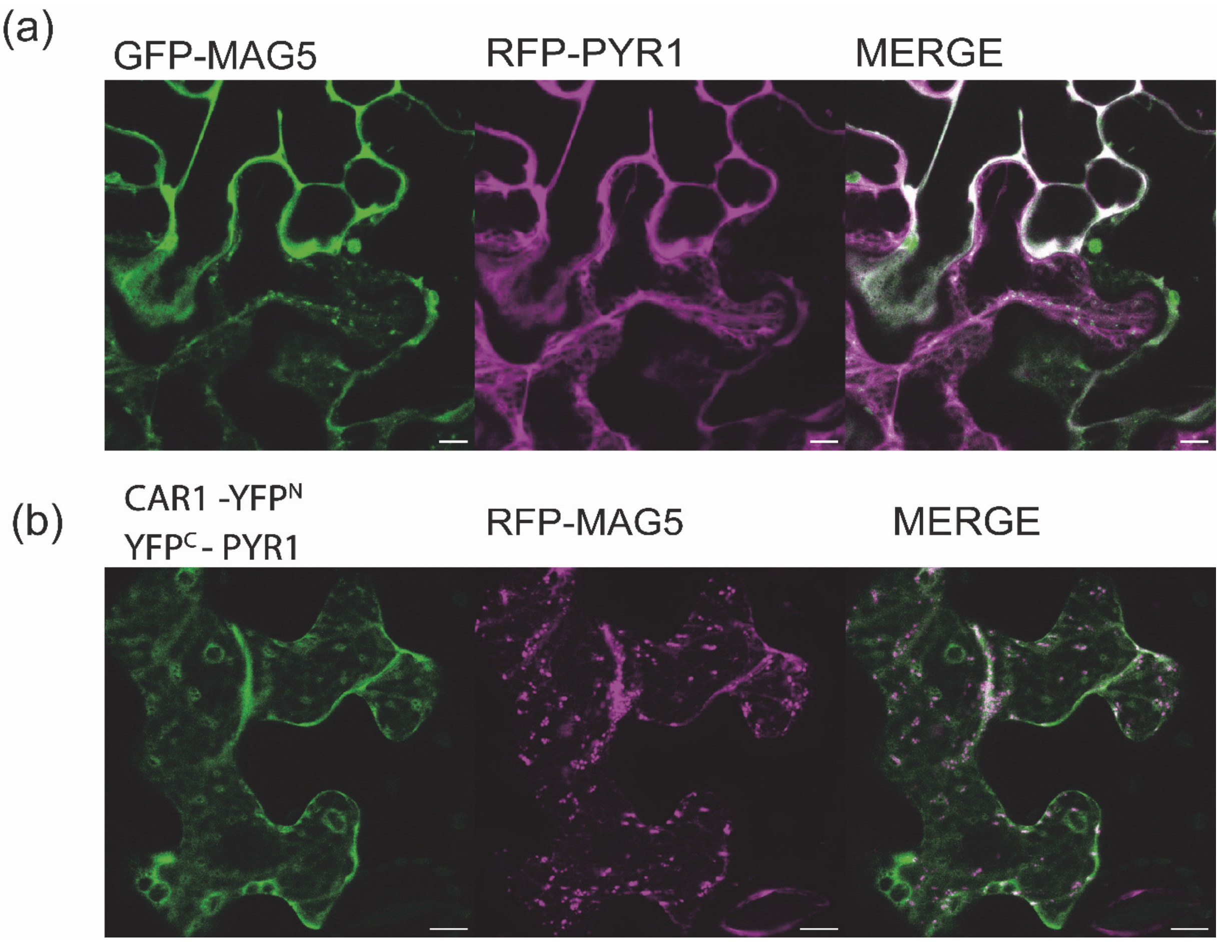


**Figure S3.** Additional evidence supporting the association of ABA receptors with MAG5-positive early secretory compartments. (a) Representative confocal images of *N. benthamiana* epidermal cells transiently co-expressing GFP-MAG5 and RFP-PYR1. (b) Representative BiFC images following transient co-expression of CAR1-YFP$^{N}$ and YFP$^{C}$-PYR1 together with RFP-MAG5. Scale bars, 10 µm.

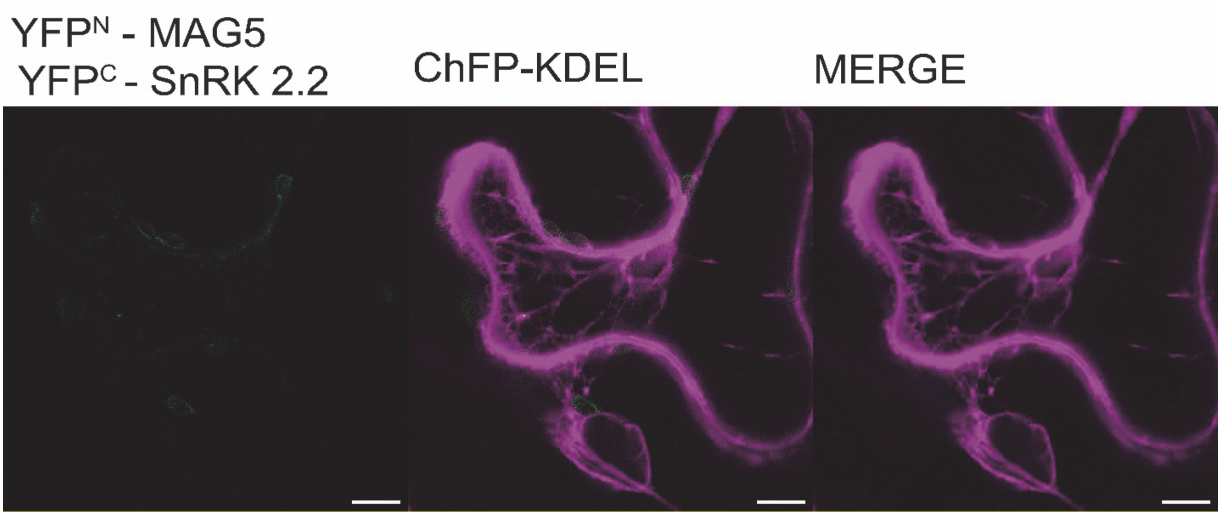


**Figure S4.** Negative controls for the BiFC assays shown in Fig. **2c**. Representative BiFC images following transient co-expression in *N. benthamiana* epidermal cells of $YFP^N$-MAG5 with $YFP^c$-SnRK2.2 together with ChFP-KDEL. Scale bars, 10 µm.

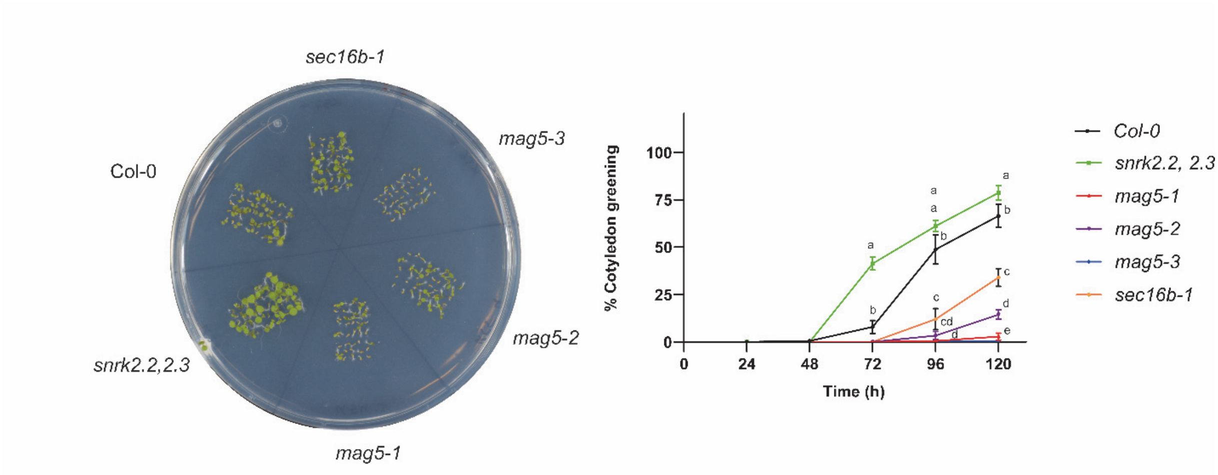


**Figure S5.** ABA sensitivity of additional *sec16* mutant alleles during cotyledon greening. Representative images of early development and quantification of cotyledon greening rates in wild-type (Col-0), the ABA-insensitive mutant *snrk2.2 snrk2.3*, *mag5-1*, *mag5-2*, *mag5-3* and *sec16b-1* seedlings grown on ½ MS control medium or supplemented with 0.3 µM ABA. Cotyledon greening was scored at the indicated time points. Data represent the mean ± SEM (n=3). Different letters indicate statistically significant differences for each time point ($P < 0.05$, one-way ANOVA with Tukey HSD test).

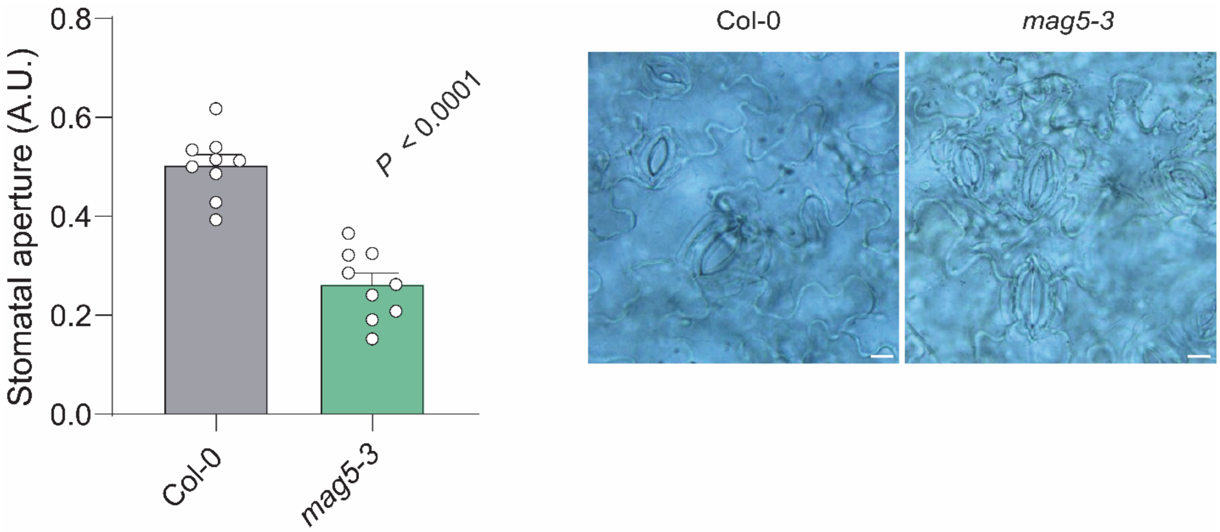


**Figure S6. Stomatal aperture is reduced in mag5-3 seedlings.** Stomatal aperture was quantified in cotyledons of 8-day-old Col-0 and *mag5-3* seedlings. Data are presented as individual seedlings and mean ± SEM (n = 9 seedlings). P-values (*P*) denote statistically significant differences from wild-type (Col-0) (Student's t-test). Representative images of cotyledon stomata are shown on the right. Scale bars, 10 µm.

**Table S1.** List of the *A. thaliana* lines used in this work

| ***Arabidopsis thaliana* line** | **Genetic background** | **Source** |
|---|---|---|
| *mag5-1* | Deletion causing a frameshift and a stop codon | Takagi *et al.*, 2013 |
| *mag5-2* | *WiscDsLoxHS202_06C* | Takagi *et al.*, 2013 |
| *mag5-3* | *GABI_147G11* | Takagi *et al.*, 2013 |
| *sec16b-1* | *SALK_070608* | Takagi *et al.*, 2013 |
| *snrk2.2 snrk2.3* | *GABI-Kat 807G04, SALK_107315* | Fujii *et al.*, 2007 |
| *hab1-1 abi1-2* | *SALK_002104, SALK_072009* | Saez *et al.*, 2006 |
| *pPYR1::PYR1-GFP* | *Col-0* | Moulinier-Anzola *et al.*, 2024 |
| *pPYR1::PYR1-GFP, 35S::RFP-p24δ5* | *Col-0* | This work |
| *pPYR1::PYR1-GFP, 35S::ManI-RFP* | *Col-0* | This work |
| *35S::GFP-PYR1* | *Col-0* | This work |
| *35S::GFP-PYR1 mag5-3* | *GABI_147G11* | This work |
| *35S::GFP-PYL4* | *Col-0* | Belda-Palazón *et al.*, 2016 |
| *35S::GFP-PYL4, 35S::RFP-p24δ5* | *Col-0* | This work |
| *35S::GFP-PYL4, 35S::ManI-RFP* | *Col-0* | This work |
| *35S::GFP-PYL4 mag5-3* | *GABI_147G11* | This work |
| *pMAG5::MAG5-GFP* | *Col-0* | Takagi *et al.*, 2013 |
| *pMAG5::MAG5-GFP, 35S::RFP-PYR1* | *Col-0* | This work |

**Table S2**. Constructions used in this work, describing the corresponding proteins expressed from each binary vector.

| ***Protein expression/vector*** | **Source** |
|---|---|
| $YFP^{N}$-MAG5 (pYFN43) | This work |
| RFP-MAG5 (pGWB455) | This work |
| GFP-MAG5 (pMDC43) | This work |
| GFP-PYR1 (pMDC43) | This work |
| RFP-PYR1 (pB7WGR2) | Moulinier-Anzola *et al.,* 2024 |
| $YFP^{C}$-PYR1 (pYFC43) | Belda-Palazón *et al.*, 2016 |
| $YFP^{C}$-PYL4 (pYFC43) | Belda-Palazón *et al.*, 2016 |
| ChFP-KDEL (pMOG800) | Serra-Soriano *et al.*, 2014 |
| ManI-RFP (pFCG derivative) | Bernat-Silvestre *et al.*, 2022 |
| RFP-p24δ5 (pCHF3) | Bernat-Silvestre *et al.*, 2022 |
| CAR1-$YFP^{N}$ (pSPYNE) | Diaz *et al.*, 2016 |
| $CAR1^{DADA}$-$YFP^{N}$ (pSPYNE) | Diaz *et al.*, 2016 |
| $YFP^{C}$-SnRK2.2 (pYFC43) | Peirats-Llobet *et al.*, 2016 |

**Table S3.** List of primers used in this work.

| Name | Sequence | Application |
|---|---|---|
| MAG5_CDS_Fw_attB1 | GGGGACAAGTTTGTACAAAAAAGCAG GCTTCATGGCTTCGACTGCTGATTTC | Clonning into pDONR™/Zeo |
| MAG5_CDS_Rv_attB2 | GGGGACCACTTTGTACAAGAAAGCTGG GTATTACAGTTCAACTTCCTGAAG | Clonning into pDONR™/Zeo |
| PYR1_qPCR_Fw | GTGCATCGGTTCGAG | RT-qPCR analysis |
| PYR1_qPCR_Rv | CACCTGAGAACCACTTCC | RT-qPCR analysis |
| PYL4_qPCR_Fw | CTCCGGGACCGTCGTTGT | RT-qPCR analysis |
| PYL4_qPCR_Rv | GGGTGGTGAAAGCCGGAA | RT-qPCR analysis |
| RAB18_qPCR_Fw | TGGCTTGGGAGGAATGCTTCA | RT-qPCR analysis |
| RAB18_qPCR_Rv | CCATCGCTTGAGCTTGACCAGA | RT-qPCR analysis |
| RD29B_qPCR_Fw | CTTGGCACCACCGTTGGGACTA | RT-qPCR analysis |
| RD29B_qPCR_Rv | TCAGTTCCCAGAATCTTGAAC | RT-qPCR analysis |
| RD29A_qPCR_Fw | GCCGAGAAACTTCAGATTGG | RT-qPCR analysis |
| RD29A_qPCR_Rv | CCATTCCTCCTCCTCCTTTC | RT-qPCR analysis |
| ABI5_qPCR_Fw | CAATAAGAGAGGGATAGCGAACGAG | RT-qPCR analysis |
| ABI5_qPCR_Rv | CGTCCATTGCTGTCTCCTCCA | RT-qPCR analysis |
| PDF2_qPCR_Fw | TCAACATCTGGGTCTTCACTTAGC | RT-qPCR analysis |
| PDF2_qPCR_Rv | GATGCAATCTCTCATTCCGATAGTC | RT-qPCR analysis |

**Table S4.** List of antibodies used in this work.

| Name | Source | Animal | Dilution |
|---|---|---|---|
| Anti-PYR1 | Agrisera AS13 2634 | Rabbit | 1:10000 |
| Anti-PYL4 | Yu *et al.*, 2026, Molecular Plant | Rabbit | 1:1000 |
| Anti-$H^{+}$ATPase | Agrisera AS07 260 | Rabbit | 1:5000 |
| Anti-CNX1/2 | Agrisera AS12 2365 | Rabbit | 1:2500 |
| Anti-β Tubulin | Santa Cruz sc-166729 | Mouse | 1:2000 |
| Anti-rabbit IgG-HRP | Jacksons #111035144 | Goat | 1:20000 |
| Anti-mouse IgG-HRP | Jacksons #115035146 | Goat | 1:20000 |
